# Altered expression of Api5 affects breast carcinogenesis by modulating FGF2 signalling

**DOI:** 10.1101/2021.11.02.466904

**Authors:** K Abhijith, Debiprasad Panda, Radhika Malaviya, Gautami Gaidhani, Mayurika Lahiri

## Abstract

Apoptosis or programmed cell death plays a vital role in maintaining homeostasis and, therefore, is a tightly regulated process. Deregulation of apoptosis signalling can favour carcinogenesis. Apoptosis inhibitor 5 (Api5), an inhibitor of apoptosis, is upregulated in cancers. Interestingly, Api5 is shown to regulate both apoptosis and cell proliferation. To address the precise functional significance of Api5 in carcinogenesis here we investigate the role of Api5 in breast carcinogenesis.

Consistently, *in-silico* analysis revealed elevated levels of Api5 transcript in breast cancer patients which correlated with poor prognosis. Overexpression of Api5 in non-tumorigenic breast acinar cultures resulted in increased proliferation and cells exhibited a partial EMT-like phenotype with higher migratory potential and disruption in cell polarity. Furthermore, during acini development, the influence of Api5 is mediated via the combined action of FGF2 activated PDK1-Akt/cMYC signalling and Ras-ERK pathways. Conversely, Api5 knock-down downregulated FGF2 signalling leading to reduced proliferation and diminished *in vivo* tumorigenic potential of the breast cancer cells. Thus, taken together, our study identifies Api5 as a central player involved in regulating multiple events during breast carcinogenesis.

## Introduction

Apoptosis Inhibitor 5 (Api5), also known as Aac11 (Anti-apoptotic clone 11)/ MIG 8(Migration Inducing Gene 8) and FIF (FGF2 interacting factor), is a 55 kDa protein localised in the nucleus(Tewari et al. 1997). The protein has a Leucine Zipper Domain (LZD), an LxxLL motif and a nuclear localisation signal (NLS)(Han et al. 2012). Tiwari *et al.* discovered Api5 as a cDNA clone that helped in the survival of cells upon serum deprivation(Tewari et al. 1997). Following this study there have been several studies to understand the function and regulation of the protein. In 2006, Morris *et al.* identified the role of Aac-11 in the transcriptional regulation of E2F1, where they demonstrated Aac-11 to inhibit E2F1-mediated apoptosis (Morris et al. 2006). Interestingly, Navarro *et al.* in 2013 reported Api5, a mammalian homolog of Aac-11, to regulate the expression of several genes required for G1/S transition that are under the control of E2F1 (Garcia-Jove Navarro et al. 2013). Together the data confirmed that Api5 could regulate both apoptosis and proliferation through E2F1. Further studies mainly focused on the regulation of apoptosis by Api5 such as its interaction and inhibition of Caspase-2 (Imre et al. 2017) and activation of ERK-mediated Bim degradation through FGF2 (Noh et al. 2014). A recent study from our lab has identified the regulators of Api5 acetylation and its’ requirement during the cell cycle (Sharma and Lahiri 2021).

Early reports started pointing at the possible tumour promoting role of Api5. In 2000, Kim et al. demonstrated that Aac11 overexpression could protect human cervical cancer cells from apoptosis (Kim et al. 2000), followed by a report on non-small cell lung cancer (NSCLC) and breast cancer. In NSCLC, higher levels of API5 were associated with poor survival of patients (Sasaki et al. 2001), while Api5 expression levels were observed to be elevated in Tamoxifen (a hormonal therapy drug that binds to the oestrogen receptor)-resistant breast carcinomas (Jansen et al. 2005). Cho *et al.* reported higher expression of Api5 in cervical cancer tissues as well (Cho et al. 2014). A recent report by Basset et al. shed more light on the role of Api5 on breast cancer, where they reported that Api5 interacted with oestrogen receptor alpha (ERα) through the LxxLL motif. They also showed reduced *in vivo* tumorigenicity upon knock-down of Api5 in xenografted MCF7 cells in mice (Basset et al. 2017).

Interestingly, Di Benedetto and her group reported that Api5 inhibition reduced angiogenesis and elevated apoptosis in xenograft models, thereby highlighting the potential use of Api5 as a therapeutic target in metastatic breast cancers that are resistant to chemotherapy (Bousquet et al. 2019). These reports suggested that Api5 could be functioning as a tumour promoter by regulating various signalling mechanisms. However, further investigations are required for establishing the role and necessity of this protein during carcinogenesis. Investigating the role of Api5 in breast cancer could be interesting as earlier reports do suggest a strong involvement of this protein in breast carcinogenesis.

Moreover, breast cancer is also a leading cause of mortality in women worldwide. GLOBOCAN 2020 data reports that breast cancer-associated mortality to incidence ratio is higher in developing countries like India. Continuous research and development have paved the way for better diagnosis and targeted therapy for breast cancer; however, taming the increase in incidence and mortality is still a challenging problem that needs to be attended to. Studying the function of Api5 in breast carcinogenesis could provide a possible biomarker or drug target for better management of the disease.

In our study, we report the role of Api5 in breast carcinogenesis using multiple models by altering the level of Api5 protein expression. Altered expression of Api5 affected proliferation, apoptosis, cell polarity and cell migration in both non-tumorigenic and tumorigenic cell lines. Api5 overexpression in MCF10A breast epithelial cells activated FGF2 signalling, leading to PDK1-Akt/cMYC activation during the early days of acinar morphogenesis. Activation of this led to elevated proliferation, migration, partial EMT and loss of polarity as was observed in Api5 overexpressed MCF10A cells. 3D cultures of these cells showed altered morphological changes and reduced apoptosis, which is speculated to be functioning through the FGF2-mediated activation of ERK signalling that was observed in the later days of acinar growth. Interestingly, reduced levels of Api5 resulted in the lower tumorigenic potential of the malignant breast cancer cell line, MCF10CA1a and impeded FGF2 signalling. Our findings provide insights into the requirement of Api5 and how it regulates several cellular processes that, upon deregulation, can lead to breast carcinogenesis.

## Results

### Api5 transcript levels are up-regulated in breast cancer and is associated with poor patient survival

To investigate the expression pattern of *API5* in breast cancers, *in silico* analysis was performed to compare *API5* expression between normal and tumour tissues of the breast using the GENT2 database. GENT2 is an online tool containing gene expression data from different cancers and compares it to normal tissues(Park et al. 2019). The log2 gene expression plot shows that Api5 transcript levels are up-regulated in breast cancer tissue samples when compared to normal breast tissues (Figure 1A). Molecular subtyping of breast cancer helps to classify breast cancer patients based on the Pam50 analysis or hormone receptor, proliferation marker and other factors. This subtyping also helps in creating treatment regimens and targeted therapies. Based on the subtyping, breast cancer can be divided into luminal A, luminal B, Her2-enriched and basal subtypes(Guiu et al. 2012). The basal subtype comprises mostly triple-negative breast cancers (TNBC) and presents the most difficult to manage breast cancer population. On comparing the *API5* transcript expression between the normal breast tissue and the various molecular subtypes of breast cancer, expression of *API5* was observed in all the subtypes (Figure 1B). Further Kaplan-Meier survival analysis of the survival probability of a patient with high / low expression of Api5 (divided by median cut-off) demonstrated poor survival of patients with high Api5 expression (Figure 1C). Both these analyses were performed using data from the GENT2 database.

**Figure 1:**
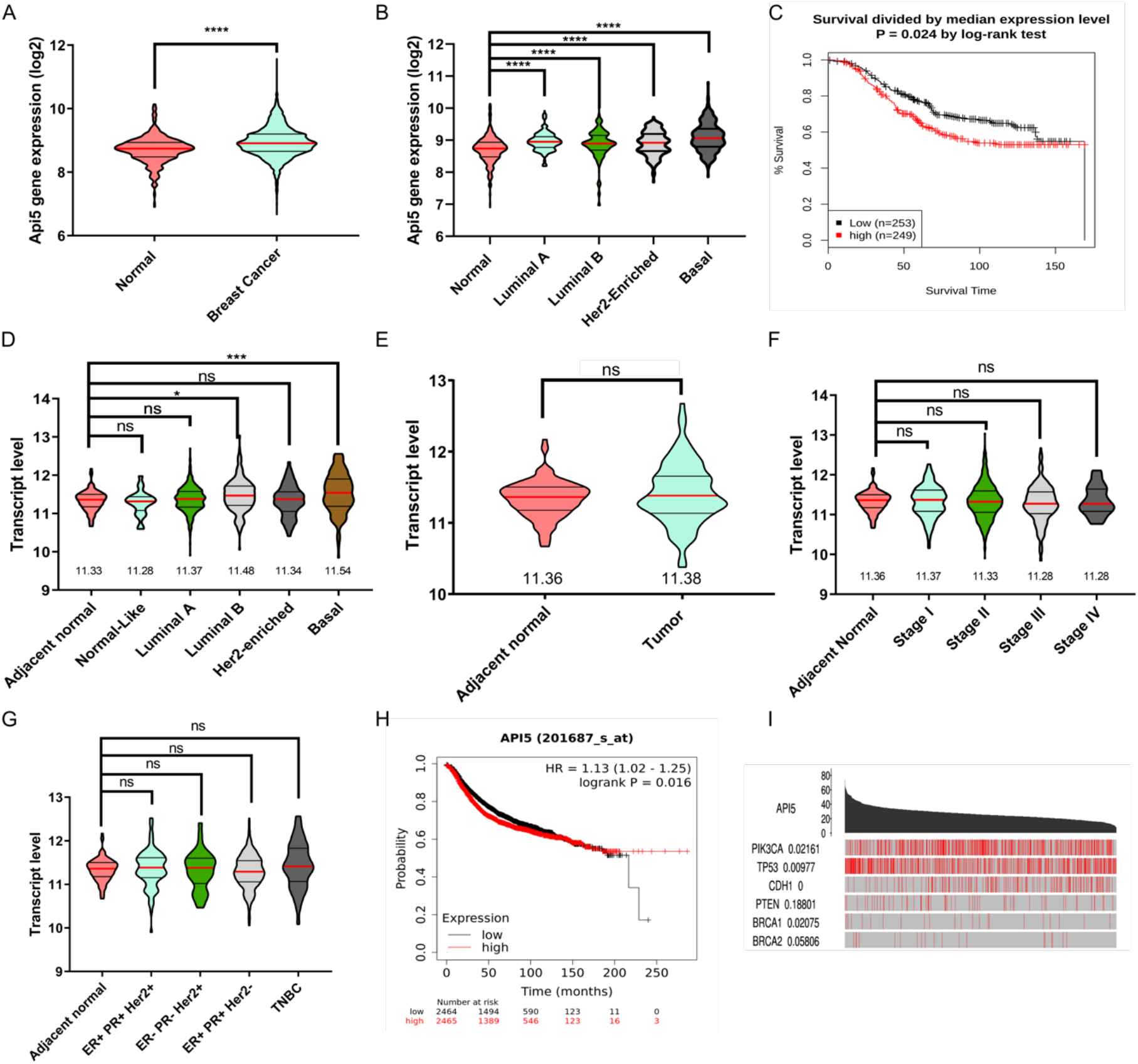
Api5 transcript levels are up-regulated in breast cancer and is associated with poor patient survival. (A) Transcript expression data (log2) of Api5 obtained from the GENT2 database is plotted. Significantly higher expression of Api5 is observed in cancer samples than in normal. Statistical analysis was performed using the Mann-Whitney test. *P<0.05, **P<0.01, ***P<0.001and ****P<0.0001. (B) API5 expression data downloaded from GENT2 was used to compare the expression across different subtypes in comparison to the adjacent normal. Statistical analysis was performed using the Kruskal-Wallis test followed by Dunn’s post hoc test. *P<0.05, **P<0.01, ***P<0.001and ****P<0.0001. (C) Kaplan Meier plot showing survival probability of breast cancer patients divided into high or low API5 expression (Median cutoff). Black line indicate survival probability of low API5 expressing patients, and red indicates high API5 expression. Significance test data is provided by the online tool. *P<0.05. (D) API5 transcript level expression obtained from TCGA database and manually analysed using Graph Pad Prism and plotted. Expression of API5 across different molecular subtype was compared to the adjacent normal sample. Transcript expression of API5 from TCGA database compared across (E) adjacent normal and tumour tissues, (F) different stages of breast cancer, and (G) receptor status-based subtypes. Statistical analysis performed using the Kruskal-Wallis test followed by Dunn’s post hoc test. *P<0.05, **P<0.01, ***P<0.001and ****P<0.0001. (The results published here are in whole or part based upon data generated by the TCGA Research Network: https://www.cancer.gov/tcga). (H) Kaplan Meier plot showing the probability of patient survival compared between API5 mRNA high or low samples. Data obtained from online tool kmplot. Expression data is divided based on the median value of the expression. Black points show low API5 expression while red show high API5 expression. The significance test was carried out by the online tool and displayed as provided. (I) API5 expression and occurrence of mutations in genes were compared and plotted using the TCGA portal online tool. Significance test was performed using the online tool and displayed as provided.

When Api5 transcript levels were compared across the different molecular subtypes of breast cancer using data from the TCGA database, the basal subtype showed higher expression of *API5* when compared to the other molecular subtypes and adjacent normal tissues (Figure 1D). However, no significant changes were observed when Api5 transcript levels were compared between adjacent normal and the hormone-receptor based subtypes as well as with the various stages of breast cancer (Figure 1E-G).

To validate Api5 mRNA expression-based patient survival prediction, a Kaplan-Meier plot was generated using KM-plotter (Nagy et al. 2021). KM-plotter uses data from multiple online databases including GEO (NCBI). Using Jetset best probe for *API5* expression and median cut-off for high versus low, the plot suggested that higher *API5* expression is associated with poor breast cancer patient survival (Figure 1H). Using TCGAportal (Xu et al. 2019) online tool, we found higher Api5 expression is associated with mutations in PIK3CA (PI3-kinase catalytic subunit alpha), TP53 and CDH1 (E-cadherin), suggesting the possible deregulation of critical pathways associated with high API5 levels (Figure 1I).

Furthermore, immunohistochemical analyses on breast cancer tissue samples were carried out to investigate Api5 expression pattern in Indian breast cancer patients. The expression levels were assayed by visual quantification followed by H-score calculation. When compared with adjacent normal or reduction mammoplasty tissues, tumour tissues showed significantly higher Api5 expression. Also, Api5 expression levels were high in Stage 2 breast cancer samples compared to Stage 1. However, there was no difference in Api5 expression between ER+ and TNBC subtypes (Supplementary Figure S1A-D).

Data from the patient samples confirm that Api5 plays a significant role in breast cancer. Elevated expression of Api5 in tumour tissues further supports the possibility that Api5 is a tumour promoter in breast malignancies. To investigate this further, we decided to overexpress Api5 in a non-tumorigenic breast epithelial cell line and study its effects on acinar phenotypes.

### Api5 overexpression in non-tumorigenic breast epithelial cells alter acinar morphogenesis due to increase in proliferation

MCF10A is a non-tumorigenic breast epithelial cell line that form growth-arrested acinar cultures when cultured in laminin-rich extracellular matrix (Matrigel^®^). This is a well-established model to study the transformation of breast epithelial cells in response to tumorigenic signalling (Anandi et al. 2017). To study the effect of Api5 overexpression on cellular transformation of the breast acini, Api5 was overexpressed in MCF10A cells (will be called Api5 OE henceforth) using lentiviral transduction followed by FACS sorting (Supplementary Figure S2A). Following 16 days of culturing, Api5 OE cells formed acini as shown in Figure 2A. On performing morphometric analysis, it was observed that Api5 OE acini have larger surface area (Figure 2B), volume (Figure 2C), increase in number of cells per acini (Figure 2D) and filled lumen phenotype (Figure 2E). There was no difference in the sphericity of Api5 OE acini when compared to the control acini (Supplementary Figure S2B) suggesting that the overall sphericity is maintained. Similarly, Api5 overexpression did not produce protrusion-like structures in the 3D cultures (Supplementary Figure S2C). Api5 OE acini were not growth arrested by day 16 and continued to proliferate as increased Ki67 protein expression was observed when compared to the control (Figure 2F-G). More than 80% of Api5 OE acini showed greater than six Ki67 positive nuclei per acini (6 cells per acini= approx. 33% of cell/acini) as shown in Figure 2H-I. On checking for the expression of PCNA, another proliferation marker, in the Api5 OE acini, an increase in PCNA levels was observed in Api5 OE when compared to the control (Figure 2J-K), thus further confirming that Api5 overexpression leads to increased proliferation. These results indicate the possibility that overexpression of Api5 in a non-tumorigenic epithelial cell line grown as acinar cultures may result in cellular transformation. To further prove this, anchorage independent growth of Api5 OE cells dissociated from the 16-day acini was investigated. Api5 OE cells formed colonies on soft agar (Figure 2L), indicating that overexpression of Api5 leads to a transformation of non-tumorigenic breast epithelial cells grown as acinar cultures.

**Figure 2:**
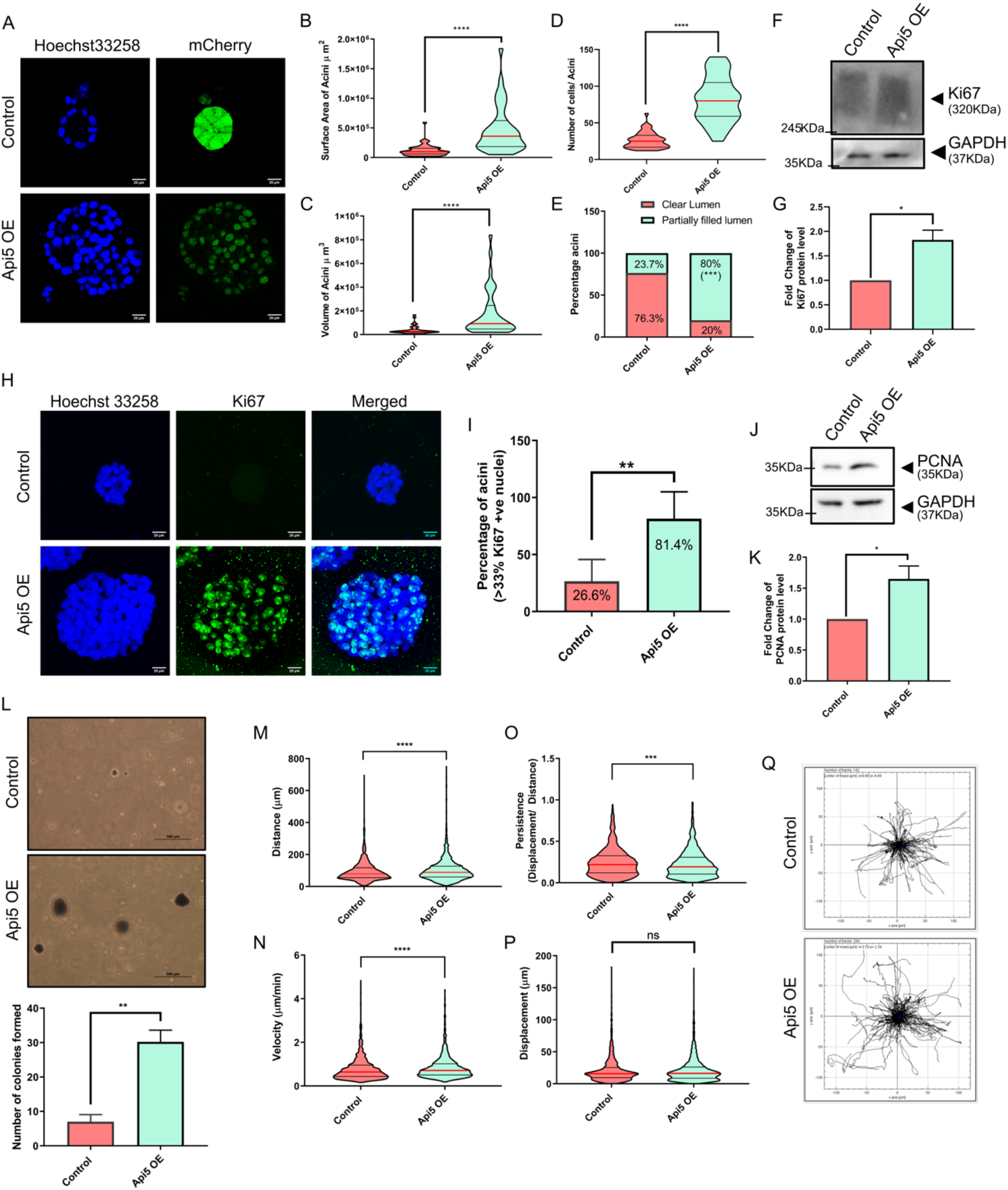
Overexpression of Api5 in MCF10A cells alter acinar morphology with increased proliferation. (A) Representative image of day 16 acini showing nuclei stained with Hoechst 33258 (blue) and mCherry (green). Violin plot showing (B) surface area and (C) volume of day 16 acini measured using Huygens software (SVI, Hilversum, Netherlands). (D) Violin plot showing number of cells in each acini, manually counted in day 16 Hoechst 33258-stained acini. Statistical analysis was performed using the Mann-Whitney test. *P<0.05, **P<0.01, ***P<0.001and ****P<0.0001. (E) Bar diagram showing percentage of acini with partially or entirely filled lumen. Day 16 acini stained with Hoechst 33258 were analysed manually by Huygens software. (F) Lysates collected from Api5 OE day 16 acini were immunoblotted for Ki67, a proliferation marker (G) Quantification showing fold change in Ki67 protein level normalised to GAPDH. Statistical analysis was performed using the paired t-test. *P<0.05, **P<0.01, ***P<0.001and ****P<0.0001. (H) Representative image of day 16 acini immunostained for Ki67 (green) and nuclei with Hoechst 33258 (blue) showing increased proliferation in Api5 OE acini when compared to control. (I) The percentage of acini with more than 33% Ki67 positive nuclei were manually analysed and plotted as a bar graph. Statistical analysis was performed using unpaired t-test. *P<0.05, **P<0.01, ***P<0.001and ****P<0.0001. Data pooled from N≥3 independent experiments. (J) Protein expression of PCNA in day 16 acinar cultures obtained using western blot and quantified in (K) showing fold change of PCNA protein levels. Statistical analysis was performed using the paired t-test. *P<0.05, **P<0.01, ***P<0.001and ****P<0.0001. (L) Representative image showing colony formed on soft agar assay stained with MTT after 21 days of seeding, manually counted and represented as a bar graph showing the total number of colonies formed on soft agar assay. Statistical analysis was performed using the Mann-Whitney test. *P<0.05, **P<0.01, ***P<0.001and ****P<0.0001. Data was pooled from n≥5 independent experiments. Control and Api5 OE cells were sparsely seeded and tracked for 3 hours to analyse (M) distance travelled (N) velocity, (O) persistence and (P) displacement. All the parameters were calculated using Fasttracks tool utilising the time-course data of the cells tracked for 3 hours. (Q) Representative data showing movement of single cells in control and Api5 OE grown as monolayer cultures, sparsely seeded and tracked for 3 hours. Statistical analysis was performed using the Mann-Whitney test. *P<0.05, **P<0.01, ***P<0.001and ****P<0.0001. Data was pooled from n≥5 independent experiments.

The transformation of epithelial cells can also lead to increased migratory potential. To assess whether overexpression of Api5 in MCF10A cells increased its migratory potential, cells dissociated from 3D acinar culture were sparsely seeded and imaged for 3 hours after they had attained their morphology. The cells were tracked using Fasttracks software (DuChez 2017) (FastTracks (https://www.mathworks.com/matlabcentral/fileexchange/66034-fasttracks), MATLAB Central File Exchange. Retrieved June 1, 2021) and parameters such as speed, displacement, distance, and persistence were calculated. Significant increase in the distance travelled (Figure 2M), velocity (Figure 2N) decrease in persistence (Figure 2O) while no change in displacement (Figure 2P) was observed in Api5 OE cells when compared to the controls. The track data was further used for plotting the cellular movement on an XY plot using ImageJ (Figure 2Q). The tracking parameters suggest that Api5 OE cells move faster and cover a longer distance than the control. Further Api5 OE cells showed lower persistence suggesting that the cells were moving in a zigzag fashion rather than taking a relatively straight direction as was observed in control cells. To understand the transformative potential of Api5, Api5 OE MCF10A cells dissociated from the acinar cultures were injected subcutaneously into the flanks of athymic mice and followed for eight weeks to check if the cells had become tumorigenic *in-vivo*. The Api5 OE cells did not form tumours (data not shown) suggesting that overexpression of Api5 is capable of partially transforming breast epithelial cells grown as acinar cultures with diminished tumorigenic potential.

Interestingly our results showed that Api5 overexpression not only led to higher proliferation but also faster migration and anchorage-independent growth. This further hints at the regulation of several cellular characteristics, such as cell polarity and cell-cell junctions.

### Overexpression of Api5 in breast epithelial results in polarity disruption

MCF10A cells cultured on Matrigel^®^ form acinar structures with a single polarised layer of epithelial cells surrounding a hollow lumen that resemble the human mammary gland acini *in vivo*. Oncogene mediated transformation is often associated with loss or disruption of polarity (Javier 2008; Halaoui and McCaffrey 2015) and is one of the hallmarks of transformation. To understand whether overexpression of Api5 can lead to disruption of polarity in the acini, Api5 OE cells were cultured for 16 days and immunostained for the different polarity markers. α6-integrin that marks the basal region of the acini (Debnath et al. 2003a) was observed to be mislocalised in 81% of the Api5 OE acini compared to the control (Figure 3A and B). Similarly, laminin V, a basement membrane marker showed loss in 82% of acini with Api5 OE (Figure 3C and D), thus, confirming that overexpression of Api5 leads to basal polarity disruption. To further understand the effect of Api5 overexpression on the other polarity components in the acinar cultures, the apical polarity marker GM130 (cis-Golgi protein) and the cell-cell junction markers E-cadherin and β-catenin were studied. 83% of the Api5 OE acini had mislocalised GM130 (Figure 3E and F), where GM130 was not observed to be apically positioned to the cell nucleus(Debnath et al. 2003a). In addition, 63% of the Api5 OE acini showed loss of E-cadherin (Figure 3G and H). Interestingly, β-catenin, another cell-cell junction marker remained unaffected in the Api5 overexpressed acinar cultures (Supplementary Figure S2D and E). Taken together, our data suggests that overexpression of Api5 leads to polarity disruption in the breast acinar cultures, which can be associated with oncogene-mediated transformation. These results indicated that Api5 overexpression resulted in several characteristics changes in the epithelial cell line, possibly through epithelial to mesenchymal transition (EMT).

**Figure 3:**
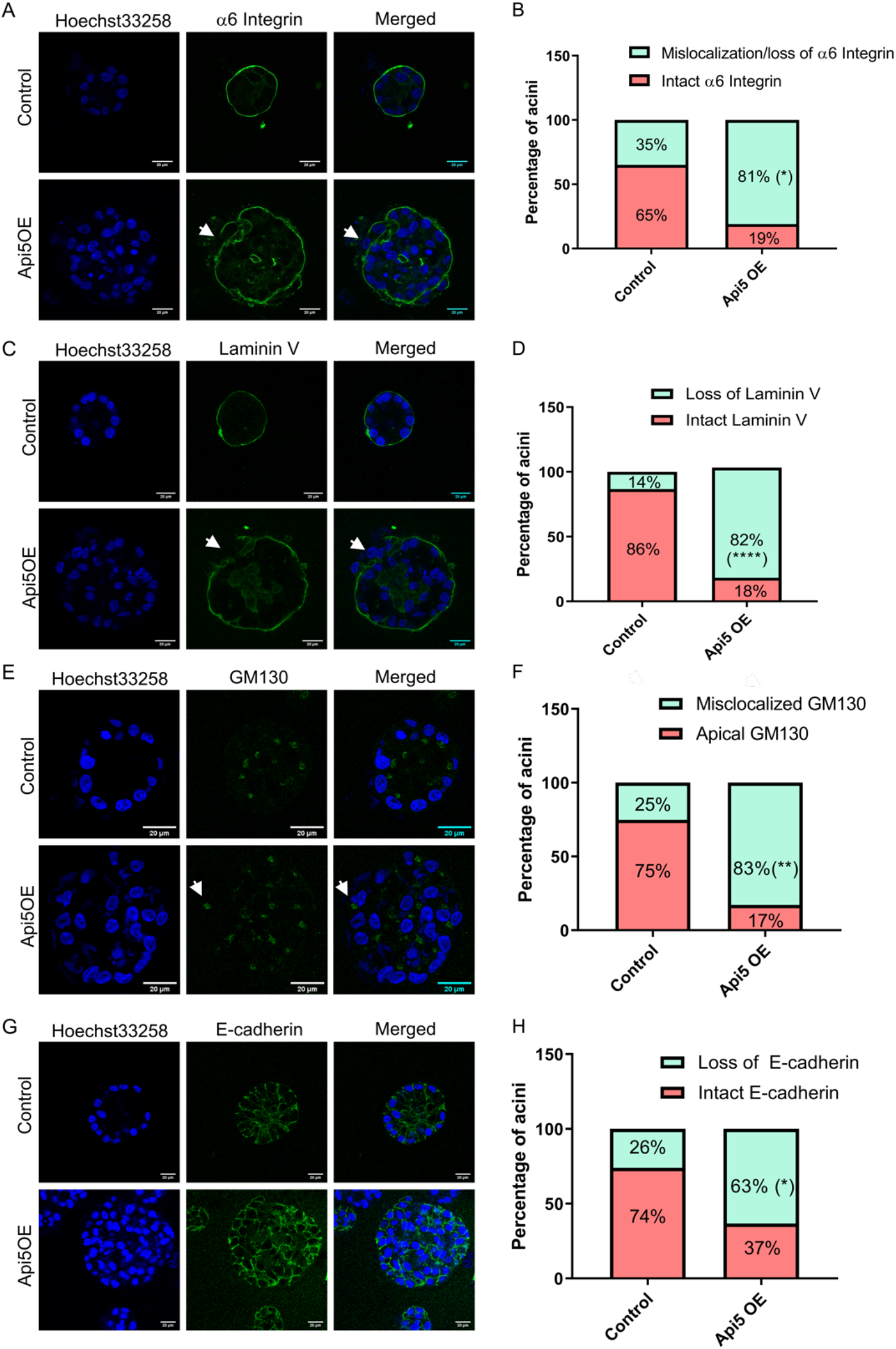
Overexpression of Api5 disrupts breast acinar polarity. (A) Representative image showing basal polarity marker, α6-integrin (green) immunostained in day 16 acini. (B) Bar diagram showing the percentage of acini with mislocalisation / loss of α6-integrin staining. (C) Representative image showing basal polarity marker Laminin V immunostaining (green) in day 16 control and Api5 OE acini. (D) Bar diagram showing the percentage of acini with mislocalisation/ loss of Laminin V staining. (E) Representative image showing GM130 (green) immunostaining in day 16 control and Api5 OE acini. (F) Quantification showing percentage of acini with mislocalised GM130 staining. (G) Representative image showing cell-cell junction marker E-cadherin (green) immunostaining in control and Api5 OE day 16 acini. (H) Bar diagram showing percentage of acini with loss of E-cadherin staining. Statistical analysis for percentage of acini was performed using unpaired t-test. *P<0.05, **P<0.01, ***P<0.001and ****P<0.0001. Data pooled from N≥3 independent experiments.

### Overexpression of Api5 in non-tumorigenic breast epithelial cells induce partial EMT-like characteristics

Transformation of breast epithelial cells are often associated with epithelial to mesenchymal transition (EMT). MCF10A cells exhibits epithelial characteristics, including expression of epithelial markers such as E-cadherin and cytokeratin. Cells acquire mesenchymal characteristics such as a change in shape and expression of mesenchymal markers such as vimentin, slug and twist upon oncogene-mediated transformation. To study whether overexpression of Api5 leads to EMT, Api5 OE cells were cultured for 16 days, lysates collected, and immunoblotting was performed. Vimentin, twist, slug and fibronectin showed 1.5-fold upregulation in Api5 OE acini when compared to the control (Figure 4A and Supplementary Figure S3A-D). However, there was no change in β-catenin and N-cadherin expression levels (Figure 4A and Supplementary Figure S3E and F). Interestingly the epithelial marker, E-cadherin showed a 2-fold increase in Api5 OE cells in comparison to the control (Figure 4A, and Supplementary Figure S3G). This was confirmed by calculating the corrected fluorescence intensity of E-cadherin in day 16 acini (Figure 4C) where a significant increase in the fluorescence intensity was observed, thereby suggesting an increase in E-cadherin levels. To further investigate whether the EMT-like phenotype that was observed upon overexpression of Api5 was a transient phenomenon or not, cells dissociated from the 16-day breast acinar cultures were grown as monolayer cultures and analysed. Similar to the phenotype observed in the acinar cultures, an increase in vimentin, slug, fibronectin and E-cadherin was observed in the Api5 OE dissociated cells (Figure 4B and Supplementary Figure S3H-K). N-cadherin expression remained unaltered in the Api5 OE dissociated cells (Figure 4B and Supplementary Figure S3L). The epithelial cell markers cytokeratin 14 and 19 both showed differential regulation in the Api5 OE dissociated cells. Cytokeratin 14 was downregulated while cytokeratin 19 was up-regulated (Figure 4B and Supplementary Figure S3M and N). The upregulation of vimentin was further confirmed when the 16-day Api5 OE breast acini as well as dissociated cells were immunostained for vimentin and increased fluorescence intensity was demonstrated for the mesenchymal marker (Figure 4D-F and Supplementary Figure S3O). Thus, taken together our data suggests that overexpression of Api5 leads to a partial EMT-like phenotype in the breast epithelial cells.

**Figure 4:**
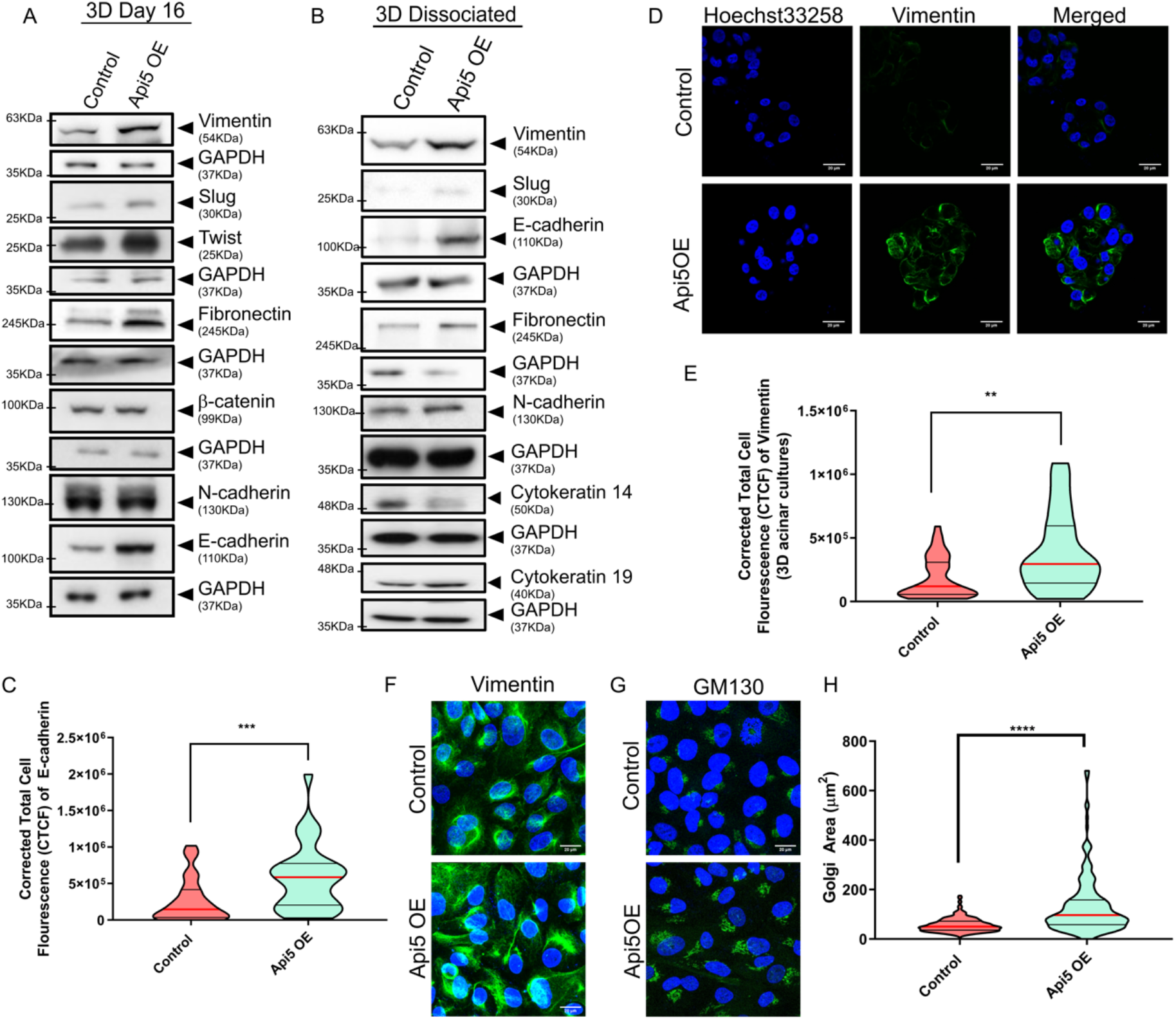
Api5 overexpression in breast acini lead to the epithelial cells acquiring partial EMT-like characteristics. Immunoblotting of lysates collected from (A) day 16 acini and (B) dissociated cells from day 16 acini and grown as monolayer cultures showing differential expression of epithelial and mesenchymal markers in Api5 OE compared to the control. (C) Violin plot showing corrected cell fluorescence of E-cadherin immunostaining of day 16 control and Api5 OE acini. (D) Representative image of Vimentin (green) immunostaining of control and Api5 OE day 16 acini. Nuclei were counterstained with Hoechst 33258 (blue). (E) Corrected cell fluorescence of vimentin quantified using ImageJ and represented as violin plot. (F) Representative image of immunostained control and Api5 dissociated cells for Vimentin (green). Nuclei was stained with Hoechst 33258 (blue). (G) Representative image of GM130 immunostained control and Api5 OE dissociated cells (GM130: green, nuclei: blue). (H) Golgi area was measured using ImageJ. Statistical analysis performed using the Mann-Whitney test. *P<0.05, **P<0.01, ***P<0.001and ****P<0.0001. Data pooled from n>5 independent experiments.

The Golgi structure has been a topic of interest in several recent studies in cancer biology. Glandular epithelial structures like breast acini have intact Golgi localised towards the apical region while more transformed or cancerous cells have dispersed Golgi (Petrosyan 2015). Anandi and group in their studies have demonstrated methylation damage leading to an aberrant Golgi phenotype in transformed MCF10A breast acinar cultures (Anandi et al. 2017). In order to investigate whether overexpression of Api5 can also lead to an aberrant Golgi morphology, Api5 OE cells were dissociated from 3D cultures and immunostained for the cis-Golgi marker GM130. Following quantification, it was observed that Api5 OE dissociated cells showed a significant increase in Golgi area compared to the control, thereby suggesting an aberrant phenotype (Figure 4G and H). Studies have reported Golgi bodies to play a major role in cell migration (Pouthas et al. 2008; Millarte and Farhan 2012; Saraste and Prydz 2019), and therefore the increase in the migratory potential along with reduced persistence that was observed in Api5 OE cells may be associated with the aberrant Golgi phenotype.

Api5 OE in MCF10A cells grown as acinar cultures resulted in significant characteristic changes in the epithelial cell line, where higher cell proliferation, loss of polarity, acquiring anchorage-independent growth and attaining a partial EMT-like phenotype were observed. Overexpression of Api5 thus resulted in the transformation of these cells.

### Reduced expression of Api5 in premalignant and malignant breast cancer cells lead to partial reversal of cancerous phenotypes

Since overexpression of Api5 was leading to a transformed phenotype, we wanted to explore whether down-regulation of Api5 could alter the cancerous phenotype in pre-malignant and malignant breast cancer cells. On investigating the protein expression of Api5 in the MCF10 cell line series (Soule et al. 1990; Dawson et al. 1996; Santner et al. 2001; Imbalzano et al. 2009), Api5 protein expression was observed to be up-regulated in MCF10CA1a cells in comparison to the non-tumorigenic MCF10A or pre-malignant MCF10AT1 cells (Figure 5A-B). shApi5 knock-down stable cells were prepared in both MCF10AT1 and MCF10CA1a and FACS-sorted (Supplementary Figure S4A and G). When Api5 KD MCF10CA1a cells were cultured for 8 days, they formed smaller spheroids as was observed by a significant reduction in surface area and volume when compared to MCF10CA1a control (Figure 5C-E). This reduction in size was further corroborated with a significant reduction in the number of cells forming the Api5 KD MCF10CA1a spheroids. 75% of the Api5 KD MCF10CA1a spheroids had structures composed of less than 100 cells (Figure 5C and F). Since overexpression of Api5 showed increased proliferation, it was intriguing to investigate whether knock-down of Api5 in the malignant cells could lead to reduced proliferation. 58% of Api5 KD MCF10CA1a spheroids showed a reduction in proliferation as was analysed by less than 50% cells per spheroid being Ki67 positive (Figure 5G and Supplementary Figure S4B). The levels of PCNA was also reduced upon knock down of Api5 (Figure 5H and Supplementary Figure S4C), thus further corroborating the observation that knock down of Api5 in malignant breast cells results in a reduction in proliferation. Furthermore, Api5 KD MCF10CA1a cells formed fewer colonies on soft agar thereby indicating that a reduction in Api5 expression negatively affected anchorage independent growth (Figure 5I and Supplementary Figure S4D). When these Api5 KD MCF10CA1a cells were injected into the flanks of athymic mice they formed tumours, although the tumour volume was significantly reduced compared to the control MCF10CA1a cells. Tumours were measured using a vernier calliper every alternative week for 8 weeks. Upon quantification, it was observed that while the MCF10CA1a control tumours continued to grow, Api5 KD MCF10CA1a tumours stopped growing and maintained a size similar to that of week 2 tumours (Figure 5J and K, Supplementary Figure S4E and F).

**Figure 5:**
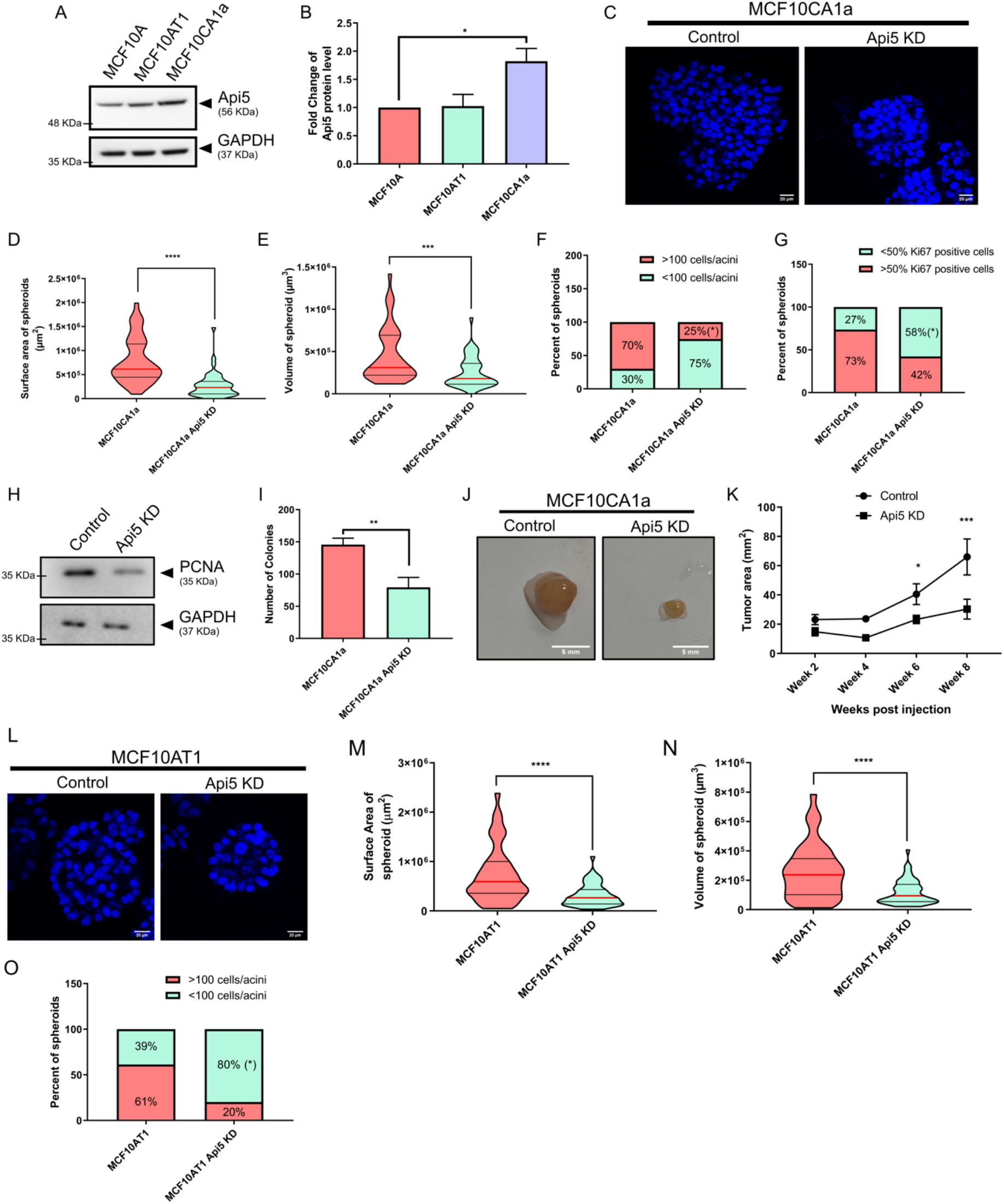
Api5 knock-down in malignant breast cells resulted in a partial reversal of cancerous phenotypes. (A) Api5 protein expression in MCF10A cell line series. (B) Quantification showing the expression levels of Api5 across the MCF10A cell line series normalised to loading control GAPDH. Statistical analysis was performed using the paired t-test. *P<0.05, **P<0.01, ***P<0.001and ****P<0.0001. Data pooled from N≥3 independent experiments. (C) Api5 KD and control MCF10CA1a cells cultured on Matrigel^®^ for seven days stained with nuclear stain Hoechst 33258 (blue). MCF10CA1a control and MCF10CA1a Api5 KD cells were immunostained with phalloidin after 7 days of culture on Matrigel^®^, and morphometric analysis was performed to calculate (D) surface area and (E) volume of spheroids using Huygens software (SVI, Hilversum, Netherlands). Statistical analysis was performed using the Mann-Whitney test. *P<0.05, **P<0.01, ***P<0.001and ****P<0.0001. Data pooled from N≥3 independent experiments. (F) The percentage of cells with >100 or <100 cells per spheroids were calculated and plotted. (G) Percentage of spheroids with more than 50% Ki67 positive cells per spheroid. Statistical analysis was performed using unpaired t-test. *P<0.05, **P<0.01, ***P<0.001and ****P<0.0001. Data pooled from N≥3 independent experiments. (H) Immunoblotting of lysates collected from day 7 3D culture probed for PCNA. (I) Plot showing total number of MTT-stained colonies formed on soft agar assay that were manually counted. Statistical analysis was performed using the Mann-Whitney test. *P<0.05, **P<0.01, ***P<0.001and ****P<0.0001. Data pooled from n>5 independent experiments. (J) Representative image of tumour dissected from flanks of athymic mice, 8 weeks post subcutaneous injection. The left panel shows MCF10CA1a control cells, while the right panel shows Api5 KD MCF10CA1a. (K) Graph showing tumour area as measured using vernier calliper at the mentioned time. For area calculation, the length of the longer axis and shorter axis were multiplied. Statistical analysis was performed using 2-way Anova followed by Sidak’s multiple comparison test. *P<0.05, **P<0.01, ***P<0.001and ****P<0.0001. Data are pooled from n>5 independent experiments. (L) Api5 KD and control MCF10AT1 cells cultured on Matrigel^®^ for 7 days and stained with Hoechst 33258. Morphometric analysis was performed on MCF10AT1 control and Api5 KD cells stained with phalloidin after 7 days of culture on Matrigel^®^, and (M) surface area and (N) volume of spheroid was measured using Huygens software. Statistical analysis was performed using the Mann-Whitney test. *P<0.05, **P<0.01, ***P<0.001and ****P<0.0001. Data pooled from N≥3 independent experiments. (O) The number of cells per spheroids was counted based on nuclear staining and plotted for percentage of spheroids with >100 (red) and <100 cells (green). Statistical analysis was performed using unpaired t-test. *P<0.05, **P<0.01, ***P<0.001and ****P<0.0001. Data pooled from N≥3 independent experiments.

Similarly, knock-down of Api5 in the pre-malignant cell line MCF10AT1 also led to a reduction in the surface area and volume of the spheroids (Figure 5L-N). 80% of spheroids formed by Api5 KD MCF10AT1 had less than 100 cells per spheroid, while in the MCF10AT1 control spheroids, more than 60% of the spheroids had more than 100 cells in the spheroid (Figure 5O). Interestingly, knock down of Api5 in MCF10AT1 spheroids did not affect proliferation as no difference was observed in the number of Ki67 positive cells between the MCF10AT1 control and Api5 KD MCF10AT1 spheroids (Supplementary Figure S4H-I). Similar to the results observed in Api5 KD MCF10CA1a cells, knock-down of Api5 in the MCF10AT1 cells also resulted in a reduced ability to grow on soft agar suggesting partial reversal of the malignant phenotype (Supplementary Figure S4J-K).

Thus, we could confirm that while Api5 overexpression alters epithelial characteristics and leads to transformation, knock down of Api5 partially reversed malignant phenotypes in malignant cells. These results confirm that Api5 is a major player involved in breast carcinogenesis.

### Api5 regulates FGF2-mediated Akt and ERK signalling

Our studies using non-tumorigenic breast epithelial and malignant cells confirmed that Api5 regulates several phenotypic characteristics that are altered due to transformation. To further understand the importance of Api5 in breast carcinogenesis, it is essential to delineate the molecular signalling involved. The coordinated functioning of Api5 and FGF2 has been intensely researched for a number of years. Studies have reported Api5 along with FGF2 to regulate ERK signalling thereby leading to Bim degradation(Krejci et al. 2007; Noh et al. 2014). Since Api5 OE MCF10A cells had filled lumen on day 16 and Bim, being an important player known to induce apoptosis during days 8 to12 of acinar morphogenesis, a number of apoptotic markers, including Bim, were studied during the 3D growth of MCF10A and Api5 OE MCF10A acini. Whole cell lysates were collected on days 4, 8, 12 and 16 of acinar growth. Overexpression of Api5 led to reduction in Bim, and active caspase 9 on day 12 of morphogenesis (Figure 6A-C). Since ERK-mediated Bim degradation has been reported to be through Api5 signalling (Noh et al. 2014), we checked whether overexpression of Api5 could lead to an alteration in ERK and MEK activity. Phosphorylation of ERK and MEK kinases were observed upon Api5 OE during day 12 of acinar growth (Figure 6A, D and E). Since ERK signalling was activated, we were interested in studying the effect on FGF2 expression levels upon overexpression of Api5 in the MCF10A acinar cultures. Increase in FGF2 protein expression was observed from days 4 to 12 in Api5 OE acini when compared to the control. This increase was a little over 3-fold on day 4 while it was 1.5-fold on day 12. By day 16, FGF2 was marginally lower in Api5 OE cells compared to the control (Figure 6A and F). Thus, overexpression of Api5 modulates a number of signalling pathways that play a role during acinar morphogenesis. Although an increase in FGF2 levels was observed on day 4, an increase in ERK signalling was observed only from day 12. Therefore, further investigation of other alternative pathways were performed. FGF2 is known to activate tyrosine kinase receptors that then activate the PI3K signalling cascade (Okada et al. 2019; Mossahebi-Mohammadi et al. 2020). In order to identify whether PI3K signalling is activated in Api5 OE MCF10A cells, phosphorylation of Akt at T308 and S473 residues were investigated. Complete activation of Akt requires initial phosphorylation at tyrosine 308 residue followed by activation at serine 473 residue. The enzyme PDK1 phosphorylates T308 residue on Akt. Overexpression of Api5 increased phosphorylation of Akt at T308 on days 4 and 8, while the Akt S473 phosphorylation was similar to that of the control cells (Figure 6A, G). Therefore it can be inferred that PDK1 is activated that may possibly lead to further activation of its downstream signalling molecule, cMYC. Upon probing for cMYC in these lysates, days 4 and 8 Api5 OE lysates showed significant increase in cMYC expression, which coincided with the phosphorylation of Akt at T308 residue (Figure 6A and H). Thus, the FGF2-mediated activation of PDK1-Akt/cMYC signalling during the initial days of growth supports the increased proliferation, protein synthesis and transformation of MCF10A cells while later activation of the ERK signalling cascade leads to reduced apoptosis and lumen filling. FGF2-PDK1/ ERK signalling is very often linked through RAS and a recent study has shown that Api5 interacts with KRAS(Bong et al. 2020), thus KRAS expression pattern was also studied. Interestingly, KRAS was up-regulated in Api5 OE 3D lysates from day 4 to day 12 and by day 16, KRAS levels were similar to that of the control (Figure 6A and I). Taken together, we propose Api5 to activate FGF2-mediated Ras-ERK and Akt signalling cascades.

**Figure 6:**
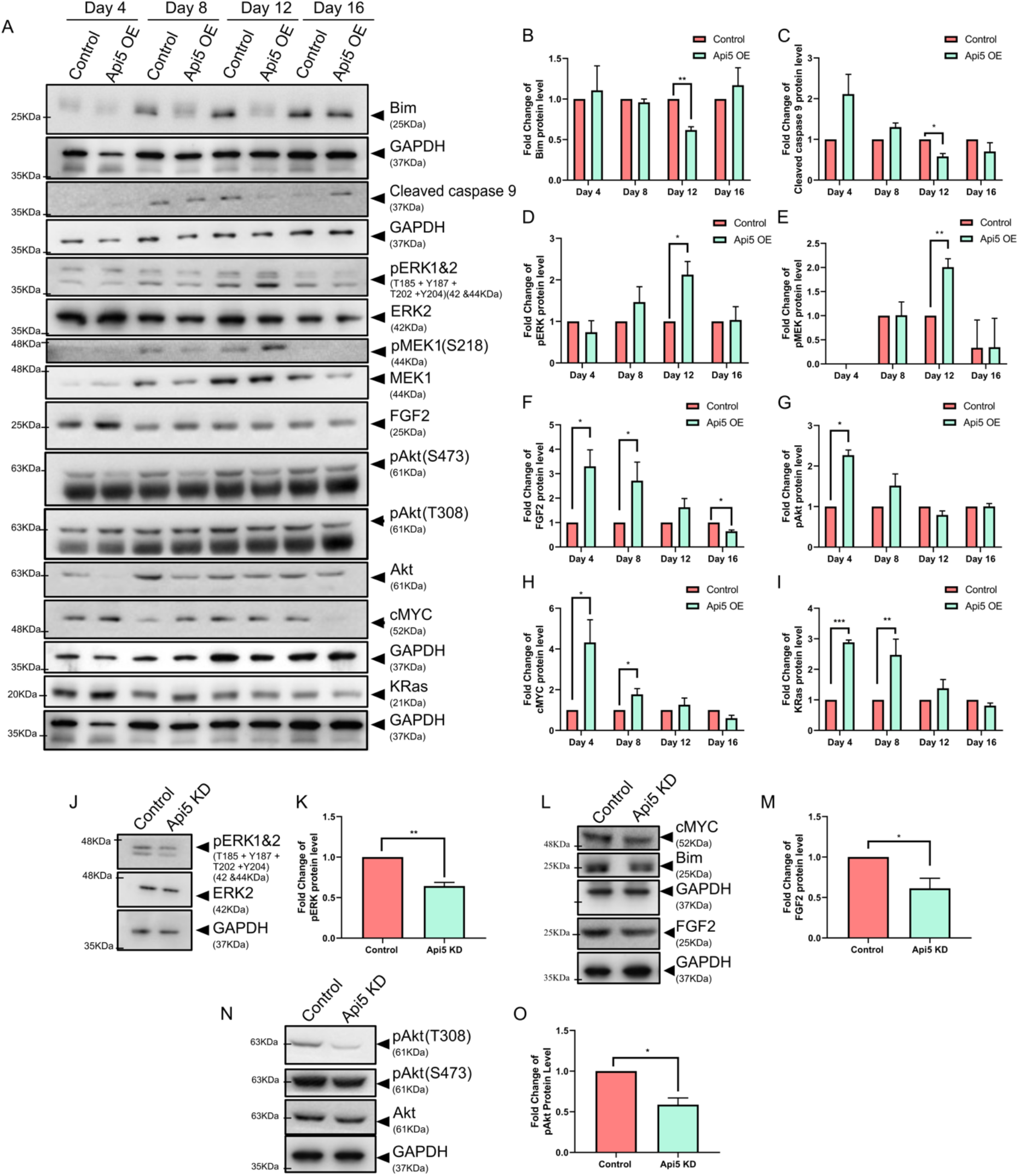
Api5 regulates PDK1/Akt and ERK pathways through FGF2. (A) Lysates were collected from control and Api5 OE acinar cultures on days 4, 8, 12 and 16 and immunoblotted to study the expression of a number of proteins. Fold change in expression between control and Api5 OE were calculated for (B) Bim, (C) cleaved caspase 9, (D) pERK 1/2, (E) pMEK1, (F) FGF2, (G) pAkt (T308), (H) cMYC and (I) KRas. Normalisation of expression was done with GAPDH for Bim, cleaved caspase 9, cMYC, and FGF2, while for pERK, pMEK and pAkt, it was with their respective total protein. (J) Immunoblotting of pERK 1&2 in day 7 MCF10CA1a control and Api5 KD spheroid cultures. (K) Quantification of the fold change in expression of pERK 1&2 normalised to total ERK2. (L) cMYC, Bim and FGF2 protein expression in Api5 KD MCF10CA1a compared to control MCF10CA1a spheroids cultured for seven days (M) Quantification of the fold-change in FGF2 expression normalised to GAPDH. (N) Lysates from day 7 MCF10CA1a control and Api5 KD spheroid cultures were immunoblotted for pAkt T308, S473 and total Akt expression levels. (O) Quantification of the fold change in expression of pAkt T308 normalised to total Akt. Data pooled from n>5 independent experiments. Statistical analysis was performed using the paired t-test. *P<0.05, **P<0.01, ***P<0.001and ****P<0.0001. Data pooled from N≥3 independent experiments.

To confirm whether Api5 can regulate the FGF2 mediated Akt and ERK signalling, Api5 KD in MCF10CA1a cells cultured on Matrigel^®^ were lysed and immunoblotted. Api5 KD resulted in a decrease in ERK phosphorylation (Figure 6J and K), suggesting Api5 may play a role in ERK signalling. Further, a reduction in FGF2 levels was also observed in Api5 KD cells compared to the control (Figure 6L and M). Interestingly, Bim and cMYC levels remained unaltered (Figure 6L, Supplementary Figure S5A and B). This may be due to the acquiring of an alternative signalling pathway when the MCF10A cells became malignant. Similarly, Akt activation was also hampered upon Api5 KD, as was demonstrated by diminished Akt phosphorylation at both S473 and T308 sites (Figure 6N and O). Western blot analysis from 3D culture lysates confirmed that Api5 played a role in both Akt and ERK signalling mediated by FGF2. Increase in cMYC expression could be responsible for the increase in the expression of mesenchymal markers. Also, growth factor-mediated signalling cascade may be providing a sustained proliferative signalling throughout morphogenesis of MCF10A, leading to partial or complete transformation of the cells even though these cells failed to form tumours in athymic mice, suggesting that there may still be other regulatory mechanism(s) playing a role. However, it is interesting to note that alteration in the expression of an anti-apoptotic protein was able to lead to an array of morphometric changes and signalling activation in the cells.

To further investigate the existence of this regulation in breast cancer patients, co-expression analysis was performed using the TCGA database. Interestingly, *BAX* and *CASPASE-9* showed a negative correlation with API5 transcript levels, but BIM mRNA levels showed a positive correlation with *API5* (Supplementary Figure S5C-E). We had observed a similar trend in the protein expression levels in Api5 OE MCF10A lysates collected on day 16. ERK2, FGF2, PDK1, KRAS and cMYC transcript levels also showed a positive correlation with API5 transcript (Supplementary Figure S5F-J) supporting the data obtained from the 3D acinar cultures. These data indicate that Api5 is regulating the FGF2-mediated PDK1 and ERK signalling in breast cancer and can possibly be used as a target for therapy.

## Discussion

Apoptosis is an important cellular process required during constant remodelling of glandular epithelium in the human breast, which is often deregulated in cancers (Mailleux et al. 2007). The role of Api5, one of the regulators in the apoptotic signalling cascade, is not well established in breast carcinogenesis. In this study, we report that deregulation of Api5 expression in breast epithelial cells leads to activation of Akt and ERK signalling pathways that affect breast morphogenesis (Figure 7).

**Figure 7:**
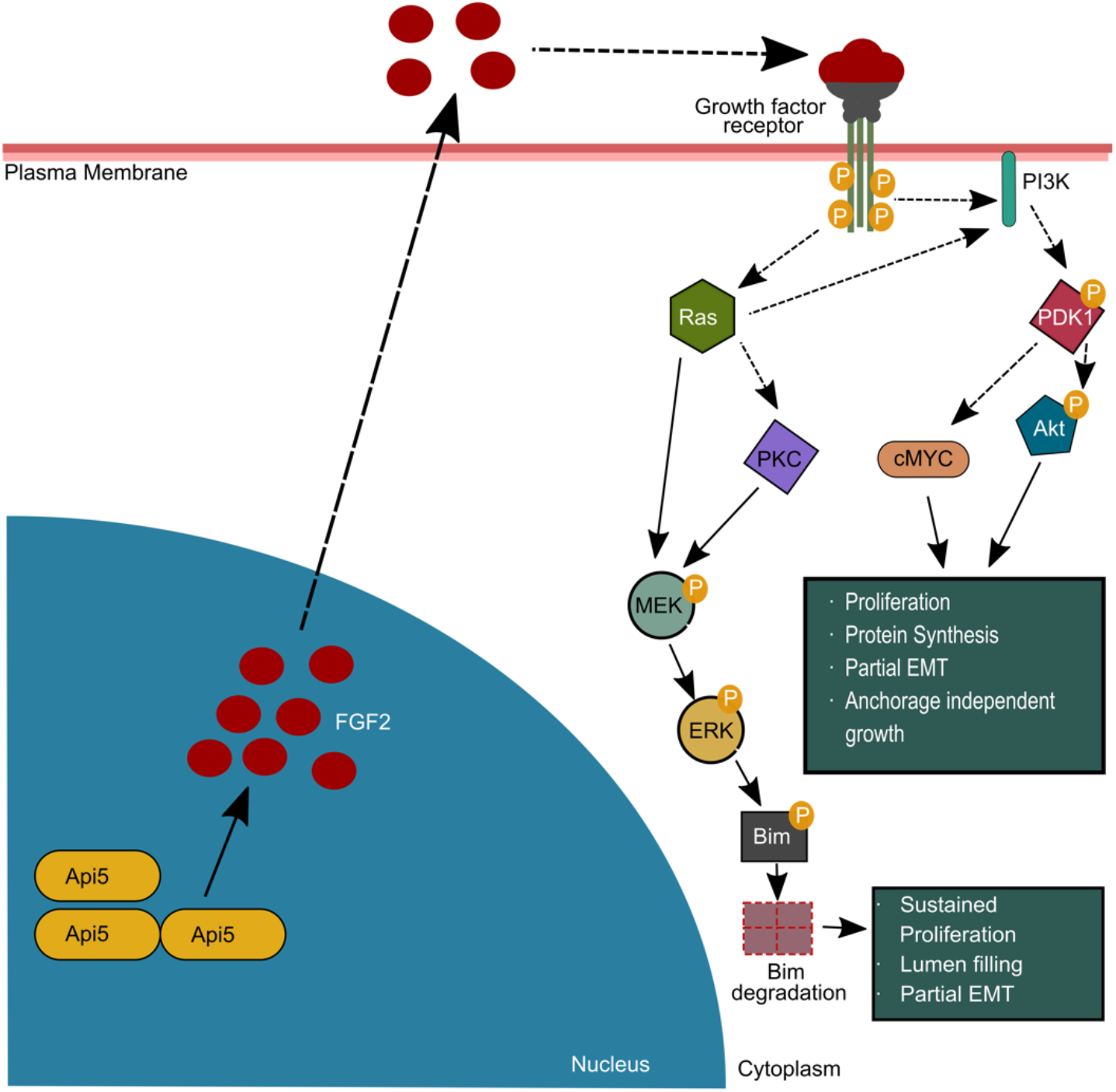
Schematic depicting the molecular mechanism of Api5-mediated regulation of PDK1/ Akt and ERK pathways, leading to the transformation of breast epithelial cells. Thick lines show signalling mechanisms revealed from experiments conducted in this study while dotted lines represents signalling mechanisms obtained from published literature. Api5 through FGF2 led to activation of growth factor receptor signalling. During the early days of morphogenesis, overexpression of Api5 led to elevated proliferation via the PDK1-Akt/cMYC pathway thus, aiding in the transformation of breast epithelial cells. Further, during the later days of morphogenesis, FGF2 signalling activated ERK-mediated Bim degradation, thereby inhibiting apoptosis and supporting sustained proliferation.

Analyses performed using multiple online tools and databases revealed that API5 transcript levels are higher in breast tumour tissues than in normal breast samples. A significant positive correlation was observed between *API5* expression and mutations in key signalling molecules such as p53 and PI3K. Mutations in these genes are known to be pathogenic and favour tumor progression (Duffy et al. 2018; Shahbandi et al. 2020; Wang et al. 2020). The data demonstrates Api5 to play an important role that favours breast tumour progression. Interestingly, recent studies have also shown that Api5 expression pattern associated with chemo-resistant TNBC patients(Bousquet et al. 2019). Furthermore, an anti-Api5 peptide, which can bind and block Api5 activity, showed anti-cancer properties and was suggested to be used as a therapeutic option (Jagot-Lacoussiere et al. 2016).

Human breast contains lobules made of numerous acini which produces milk. In each acinus, epithelial cells surround an empty lumen in which milk is secreted and is then carried through the ducts (Javed and Lteif 2013). When cultured on specific extra cellular matrices, breast epithelial cells can form similar structures *in vitro*(Swamydas et al. 2010). Such 3D acinar cultures can resemble the human mammary gland acini structurally and functionally. These spheroid cultures could provide insights into molecular events that occurs during breast carcinogenesis. MCF10A, a non-tumorigenic breast epithelial cell line, forms growth arrested acinar structures when cultured on laminin-rich extra cellular matrix (Debnath et al. 2003a). Overexpression of oncogenes such as Erbb2, Akt and cMYC led to morphometric changes in the MCF10A acini (Debnath et al. 2003b; Partanen et al. 2007). Oncogenic transformation in 3D acinar cultures can result in filled lumen, larger acinar size and formation of protrusions from the spherical structures (Debnath and Brugge 2005). These characteristic changes that epithelial cells acquire have been enumerated in the hallmarks of cancer, coined by Hanahan and Weinberg (Hanahan and Weinberg 2011) such as resisting cell death, sustained proliferative signalling, invasion and metastasis. This makes MCF10A breast acinar cultures an effective model system for studying breast acinar morphogenesis and carcinogenesis (Bessette et al. 2015; Qu et al. 2015).

In our study, MCF10A cells stably overexpressing Api5 were cultured as spheroids. Distinct differences were observed between the control and Api5 OE cells. Morphometric changes including increase in size and cell number indicated the possibility of cellular transformation, thus suggesting a plausible role of Api5 in breast cancer. Api5 OE cells showed sustained proliferative capacity even at day 16 as was demonstrated by increased levels of Ki67 and PCNA. Furthermore, knock-down of Api5 in MCF10A isogenic cell lines, MCF10CA1a (malignant) (Santner et al. 2001) and MCF10AT1 (pre-malignant) (Dawson et al. 1996) led to reduction in both the size of spheroids as well as proliferation.

MCF10A acini have epithelial cell characteristics with basal and apical polarity (Debnath et al. 2003a). Several reports have demonstrated that loss of polarity and reduced lumen size are associated with breast cancer progression (Debnath and Brugge 2005; Imbalzano et al. 2009; Halaoui et al. 2017). In our study we observed a similar phenotype in MCF10A acini overexpressing Api5. Both Laminin V and α6-integrin were mislocalised or lost at several regions in the Api5 OE MCF10A acini. Golgi, in MCF10A acini, is localised to the apical region (Gaiko-Shcherbak et al. 2015). Api5 OE led to the mis-localisation of Golgi to the basal regions.

Epithelial cell characteristics are mostly altered during transformation (Roche 2018). Epithelial cells gain mesenchymal characteristics such as the expression of proteins like Slug, Twist, and Vimentin. We observed that Api5 OE led to a partial EMT-like phenotype. A partial/hybrid EMT state is observed when a cell expresses both epithelial and mesenchymal markers (Aiello and Kang 2019). Such a state can support collective cell migration and promote circulating tumour cell movement (Jolly et al. 2015). Recent reports have demonstrated that the hybrid EMT state helps cells to attain stemness characteristics and drug resistance (Saitoh 2018), both of which are also associated with Api5-FGF2 signalling (Jang et al. 2017; Song et al. 2017). Identifying whether Api5-induced partial EMT could drive drug resistance and induce stemness characteristics in cancer cells will aid in developing treatment strategies against chemo-resistant cancers.

Increase in migratory potential is a common phenotype observed in malignant cancer cells(Paul et al. 2017). In some scenarios they follow collective cell migration while single cell migration is also observed. The single cells detached from the tumour travel to another tissue, often leading to metastasis (Blockhuys et al. 2020). We report that overexpression of Api5 led to increased single cell migration. The cells attained higher speed and travelled longer distance, however, with reduced persistence. A similar result was earlier reported when fibroblast and glioblastoma cells were treated with EGF(Kim et al. 2008). A lower persistence indicates that the directionality of cellular movement is affected. Golgi plays a major role in cell migration and directionality of movement(Millarte and Farhan 2012). Since Api5 OE resulted in the dispersal of Golgi, this may be promoting the elevated migratory potential, however, with reduced persistence(Tang et al. 2019).

Further, we also demonstrated that Api5 regulates anchorage-independent growth, a well-established property suggesting malignant transformation. Api5 OE led to anchorage-independent growth of MCF10A cells, while knock down in MCF10CA1a and MCF10AT1 showed reduced colony formation on soft agar. When the Api5 KD cells (MCF10CA1a) were injected into the flanks of mice, they formed tumours, although the size was smaller when compared to MCF10CA1a control cells. However, Api5 OE cells did not form tumours in athymic mice. This suggests that overexpression of only Api5 may not be sufficient to cause complete transformation and carcinogenesis of breast epithelial cells. A series of deregulation and genomic changes are required for a cell to be completely transformed.

During MCF10A acinar morphogenesis, cells in the lumen undergo Bim-mediated apoptosis during days 10 to 16 (Reginato et al. 2005). Api5 OE led to reduced Bim levels, thereby inhibiting apoptosis. Studies have reported that oncogenes such as Erbb2 and v-Src show a similar inhibition of Bim, thus preventing luminal cell death(Reginato et al. 2005). This reduction in apoptosis explains the presence of a partially or entirely filled lumen that was observed in the Api5 OE acini. Bim protein levels are regulated by ERK signalling where ERK activation leads to phosphorylation of Bim and thereby its degradation(O’Reilly et al. 2009). We identified that Api5 OE activated FGF2-MEK-ERK signalling during day 12 of acinar morphogenesis, which coincided with the observed Bim degradation. Knock-down of Api5 in the malignant breast cell line confirmed that Api5 can regulate the FGF2-MEK-ERK signalling cascade (Figure 7).

Api5 OE induced higher FGF2 expression from day 4 of acinar morphogenesis which activated PDK1-Akt/cMYC signalling. Knock-down of Api5 resulted in reduced activation of this signalling cascade. c-MYC expression is known to be up-regulated in several breast cancers. Higher expression is also predicted to cause poor patient outcome(Green et al. 2016). Elevated c-MYC expression also mediates EMT in breast cancers (Cho et al. 2010; Yin et al. 2017), thus possibly explaining the partial-EMT observed in Api5 OE cells. Interestingly, Partanen and group in 2007 reported that overexpression of c-MYC in MCF10A led to filling of lumen and possible transformation-like phenotypes, although interestingly, the organised and polarised acinar architecture suppressed the transforming capability of c-MYC (Partanen et al. 2007). Later Simpson *et al.* reported that endogenous and exogenous c-MYC expression are suppressed during later stages of MCF10A acinar cultures (Simpson et al. 2011). Although Api5 overexpression led to a transformed phenotype in our study, the cells did not acquire malignant potential. Interestingly, our results also indicate that Api5 overexpression induced altered signalling was restored by day 16 of acinar morphogenesis. Therefore, it is possible that acinar morphogenesis could have prevented a complete malignant transformation of overexpressed Api5 in MCF10A cells.

The ability of Api5 to regulate multiple signalling pathways may be an advantage for developing novel breast cancer treatment strategies. Chemoresistance is a complicated phenomenon that further reduces the survival chance of patients. Activation of Akt and ERK signalling are often found to be associated with chemoresistance in cancers. As mentioned earlier, several reports also implicate Api5 to play a role in chemoresistance (Jang et al. 2017; Bousquet et al. 2019). Thus, targeting Api5 might be an excellent strategy for managing such cancers.

Our investigations have uncovered a detailed understanding of the role of Api5 in breast carcinogenesis. We report that Api5 through FGF2, regulates ERK and Akt signalling, thereby leading to transformation of breast epithelial cells (Figure 7). Our data indicate Api5 to be a major regulator of several key events during breast carcinogenesis including elevated proliferation, decreased apoptosis, polarity disruption and anchorage-independent growth. This study opens up a plethora of new research opportunities as well as new possibilities for drug development.

## Materials and Methods

### Chemicals and antibodies

Cholera Toxin (C8052), Epidermal Growth Factor (E9644), Hydrocortisone (H0888), Insulin (I1882), Polybrene (H9268), Triton X-100 (T8787), Tris Base (B9754), EDTA (E6758), Na3VO4 (S6508), Protease inhibitor cocktail (P8340) and Poly-L-Lysine (P8920) were purchased from Sigma-Aldrich. Lipofectamine-2000 (11668-500) was purchased from Invitrogen, Thermo Fisher Scientific. Dispase (354235) was purchased from Corning, Sigma-Aldrich. NaF (RM1081) Na2HPO4 (GRM1417), and KH2PO4 (MB050) were purchased from HiMedia. NaCL (15918), Xylene (Q35417), Isoproopanol (Q26897), and 30% H2O2 (Q15465) were purchased from Qualigens. 16% paraformaldehyde (AA433689M) was purchased from Alfa Aesar.

Immunofluorescence staining was carried out using Ki67 (Abcam, monoclonal, ab16667), α6-integrin (Merck, monoclonal, MAB1378), Laminin V (Merck, monoclonal, MAB19562), GM130 (Abcam, polyclonal, ab30637), E-cadherin (Abcam, monoclonal, ab1416), Vimentin (Abcam, monoclonal, ab92547), and β-catenin (Abcam, monoclonal, ab32572). IHC against Api5 was performed using Api5 (Sigma polyclonal, HPA026598). Api5 (Sigma, polyclonal HPA026598 or Abnova, polyclonal, PAB7951), PCNA (Cell signalling, monoclonal, 2586), E-cadherin (BD, monoclonal, 610182), N-cadherin (Abcam, polyclonal, ab18203), GAPDH (Sigma, polyclonal, G9545), Vimentin (Abcam, monoclonal, ab92547), Slug (Cell Signalling, monoclonal, 9585), Twist (Abcam, Polyclonal, ab50581), Fibronectin (BD, monoclonal, 610077), β-catenin (BD, monoclonal, 610153), Cytokeratin 14 (Abcam, monoclonal, ab7800), Cytokeratin 19(Abcam, monoclonal, ab52625), Bim (Abcam, monoclonal, ab32158), Cleaved Caspase-9 (Abcam, polyclonal, ab2324), pERK 1&2 (Abcam, monoclonal, ab50011), ERK2 (Abcam, monoclonal, ab32081), pMEK1 (Abcam, monoclonal, ab32088), MEK1 (Abcam, monoclonal, ab32091), FGF2 (Millipore, monoclonal, 05-118), pAkt T308 (Cell Signalling, monoclonal, 4056), pAkt S473 (Invitrogen/ Biosource, monoclonal, 44-621G), Akt (Cell Signalling, monoclonal, 9272S), and cMYC (Santacruz, monoclonal, SC-40) were used for the immunoblotting experiments. Peroxidase-conjugated AffiniPure goat anti-mouse (115-035-003) and anti-rabbit(111-035-003), as well as AffiniPure F(ab′)2 fragment goat anti-mouse IgG, F(ab′)2 fragment specific (115-006-006), were obtained from Jackson Immuno Research. Hoechst 33342 (H3570), Hoechst 33258 (H3569), Alexa Fluor ^®^ 488 conjugated anti-mouse secondary antibody(A-11029), Alexa Fluor ^®^ 488 conjugated anti-rabbit secondary antibody (A-11034), Alexa Fluor ^®^ 568 conjugated anti-mouse secondary antibody(A-11004), Alexa Fluor ^®^ 568 conjugated anti-rabbit secondary antibody(A-11036), Alexa Fluor ^®^ 568 conjugated anti-rat secondary antibody(A-11077), Alexa Fluor ^®^ 633 conjugated anti-mouse secondary antibody (A-21052), Alexa Fluor ^®^ 633 conjugated anti-rabbit secondary antibody (A-21071), and Alexa Fluor ^®^ 568 phalloidin (A-12380) were bought from Invitrogen, Thermo Fisher Scientific.

### Plasmids

CSII-EF-MCS plasmid was a gift from Dr Sourav Banerjee, NBRC, Manesar, India. pCAG-HIVgp and pCMV-VSV-G-RSV-Rev plasmids were purchased from RIKEN BioResource Centre. mVenusC1 was gifted by Jennifer Lippincott-Schwartz, NIH, USA in which Api5 was cloned.

Primers used for Api5 CDS insertion to CSII-EF-MCS vector were: Forward primer: 5’-AAGGAAAAAAGCGGCCGCATATGCCGACAGTAGAGGAGCT-3’ and reverse primer: 5’-GCTCTAGATCAGTAGAGTCTTCCCCGAC - 3’.

pMD2.G and pPax2 were generous gift from Dr Manas Kumar Santra, NCCS, Pune, India. pLKO1.EGFP was a generous gift from Dr Sorab Dalal, ACTREC, Mumbai, India.

Primers used for shApi5 insertion to pLKO1 vector were: Forward primer: 5’-CCGGAAGACCTAGAACAGACCTTCACTCGAGTGAAGGTCTGTTCTAGGTCTTTTTTTG - 3’ and reverse primer: 5’-AATTCAAAAAAAGACCTAGAACAGACCTTCACTCGAGTGAAGGTCTGTTCTAGG TCTT - 3’.

### *In-silico* analyses

Using the GENT2 website, the subtype-based expression profile was plotted by selecting the subtype profile tab and providing “Api5” as gene symbol. Survival plot is presented as provided by the tool with overall survival of the patients. TCGA data (Legacy dataset) downloaded from Xena Browser (UCSC Xena) by selecting IlluminaHiSeq dataset along with clinical data. The data, merged with expression data based on sample ID using R package, is then sorted to separate adjacent normal and tumour data (Sample ID ending with “11” represents adjacent normal data while “01” represents tumour data.) Graph Pad prism (Graph Pad Software, La Jolla, CA, USA) is used for plotting the extracted data. Further Kaplan Meier analysis was carried out using the kmplot online tool. Under breast cancer, “API5” was given as a gene symbol and “201687_s at” dataset was selected and using median cut-off, the survival plot was generated. Mutation data were analysed using the TCGAportal online tool. TCGA BRCA was selected as a dataset, and Api5 was given as a gene symbol.

### Cell lines and culture conditions

The MCF10A cell line was a generous gift from Prof. Raymond C. Stevens (The Scripps Research Institute, La Jolla, CA), while MCF10AT1 and MCF10CA1a were purchased from ATCC. These cells were grown in DMEM with high glucose and without sodium pyruvate (Invitrogen) containing 5% horse serum (Invitrogen), 20 ng/ml EGF (Sigma-Aldrich), 0.5 µg/ml hydrocortisone (Sigma-Aldrich), 100 ng/ml cholera toxin (Sigma-Aldrich), 10 µg/ml insulin (Sigma-Aldrich) and 100 units/ml penicillin-streptomycin (Invitrogen). Cells were resuspended in high glucose DMEM without sodium pyruvate, containing 20% horse serum and 100 units/ml penicillin-streptomycin (Invitrogen) during passaging. The overlay medium in which the cell suspension was made for seeding (assay medium) contained DMEM without sodium pyruvate, horse serum, hydrocortisone, cholera toxin, insulin, EGF and penicillin-streptomycin. HEK 293T cell line was a generous gift from Dr Jomon Joseph (National Centre for Cell Science, Pune, India). The cells were grown in DMEM with high glucose and sodium pyruvate (Invitrogen) containing 10% foetal bovine serum (Invitrogen), and 100 units/ml penicillin-streptomycin (Invitrogen). Cells were resuspended in the same media for passaging. All cell lines were maintained in 100 mm petri dishes (Corning, Sigma-Aldrich / Nunc, Thermo Fisher Scientific / Eppendorf) at 37°C in a humidified 5% CO_2_ incubator (Eppendorf). Stable cell lines overexpressing Api5 was prepared using lentiviral-mediated transduction. Briefly, HEK293T cells were transfected with 1μg CSII-EF MCS mCherry Api5 vector having a mCherry-Api5 sequence, along with 0.5μg pCMV-VSV-G-RSV-Rev and 1μg pCAG-HIVgp for viral particle preparation using Lipofectamine 2000 (Invitrogen)-mediated transfection. Opti-MEM^®^ used for transfection was obtained from Invitrogen. DMEM containing 15% horse serum was added to the cells 24 hours post transfection. 48 hours post transfection, viral supernatant was collected and filtered through a 0.45 μm filter to remove cell debris. The viruses were then used to transduce MCF10A. 4 μg polybrene was added to the cells to increase the transduction efficiency. As control, a stable cell line expressing only mCherry was also prepared. The transduced cells were sorted using BD FACS Aria (BD Biosciences) to get a pure population with maximum number of transduced cells. Api5 KD stable cells were generated in MCF10AT1 and MCF10CA1a in a similar manner. The shRNA was cloned in pLKO.1-EGFP vector, and packaging plasmids pMD2.G and pPax2 were used for lentiviral preparation. Cells were then sorted in BD FACS Aria.

### 3D ‘on-top’ culture

3D breast acinar cultures were set up in an 8-well chamber cover glass plates (Nunc Lab-Tek, Thermo Fisher Scientific) or 12-well plates (Eppendorf) using standard protocols(Debnath et al. 2003a; Anandi et al. 2016; Anandi et al. 2017). Cultures were grown in a humidified incubator with 5% CO2 and maintained at 37°C (Eppendorf). The medium was changed every four days. For lysate collection on different days, higher cell density was seeded for Day 4 and Day 8.

For dissociation of acinar culture, Dispase™ (Corning, Sigma-Aldrich) was used. After the addition of Dispase™, the culture was incubated for 20 minutes. The dislodged acini were spun down at 900 rpm for 10 minutes, followed by 2 rounds of 1X PBS wash before plating in 12-well petri plates.

### Immunofluorescence staining

3D spheroid cultures were immune-stained with specific antibodies by established protocols(Anandi et al. 2017). For MCF10CA1a and MCF10AT1 spheroid staining, the 1X PBS and 1X IF buffer washes were added with 0.5% Triton X to aid for better penetration of antibodies. Images were captured using Leica SP8 or Zeiss LSM 710 laser scanning confocal microscope.

### Immunoblotting

Lysates from 2D or 3D cultures were collected in lysis buffer containing 50 mM Tris-HCl, pH 7.4, 0.1% Triton X-100, 5 mM EDTA, 250 mM NaCl, 50 mM NaF, 0.1 mM Na_3_VO_4_ and protease inhibitors. Immunoblotting against specific proteins was performed as per established protocols(Bodakuntla et al. 2014). The blot images represented shows all the bands that were captured using the imaging system. None of the images have been cropped. The entire blots were cut as per experimental requirement and probed for different antibodies.

### Immunohistochemistry

Formalin-fixed and paraffin-embedded breast cancer patient samples were collected from Prashanti Cancer Care Mission, Pune. The samples from each patient may contain tumour area, adjacent normal tissue, lymph nodes, and the reduction mammoplasty tissue. Tissue sectioned using Leica Microtome at 3 μm thickness were taken to poly-L-Lysine (Sigma) coated slides. Tissues are deparaffinised by overnight incubation at 62°C followed by xylene (Fisher scientific) washes. After rehydration of tissues with decreasing concentration of iso-propanol followed by water, endogenous peroxidases are blocked by incubating with 3% H2O2 (made from 30% H2O2, Fisher scientific, by diluting with methanol). Antigen retrieval was carried out using 0.001M EDTA buffer (pH 7.4). Following blocking using 3% BSA, Api5 Antibody (1:2000 dilution, Sigma HPA026598) was added on top of the tissues and incubated at room temperature for 2 hours. Further, tissue is probed with HRP conjugated secondary antibody (Dako) for 30 minutes and then developed with DAB (Dako). The nuclei are counterstained with 10% haematoxylin and mounted with DPX mountant (Fisher Scientific). The analysis was carried out by observing slides at 20X magnification of compound microscope.

H-score for IHC is calculated as follows:

H-Score = Intensity score X Percentage positivity score

The intensity score and percentage positivity score ranges from 0 to 3, and the maximum H score is 9. Three individuals calculated the scores independently by observing ten different positions on the slide corresponding to the tissue ID. The median value from this observation was plotted on the graph.

### Single Cell Migration

Cells were sparsely seeded in 8-well Lab-Tek chamber cover glass slides and incubated for 16 hours in a 37°C incubator supplied with 5% CO_2_. After staining the nuclei with Hoechst 33342 (Invitrogen), cells were imaged at 20X magnification using Leica SP8 confocal microscope for 3 hours.10 positions were selected for each well, and images were taken every 2 minutes. The data was processed using Fasttracks software (DuChez 2017) and plotted using Graph Pad Prism (Graph Pad Software, La Jolla, CA, USA).

### Soft Agar Assay

Soft agar colony formation assay was performed as detailed earlier (Anandi et al. 2017). Images were acquired using the 10X objective of a Nikon phase-contrast microscope. Ten randomly selected fields were imaged, and the colonies were manually counted. Clonogenicity assay was performed by sparsely seeding cells in 6-well plates (Corning) with three replicates for each set. After 7 days of incubation, cells were stained with crystal violet (HiMedia) and images were captured using HP Scanner after drying the plates.

### *In-vivo* tumorigenicity assay

6 × 10^6^ cells were injected subcutaneously in the flanks of athymic mice (Foxnl ^nu^ / Foxnl ^nu^, 6-8 weeks old) mixed with 1:1 diluted Matrigel^®^ PBS mixture. Tumour size was measured with a vernier calliper. Furthermore, after 8-12 weeks, mice were sacrificed, and the tumour was dissected out. The tumour was then fixed with 10% formaldehyde and embedded in paraffin wax. Tumour area was calculated by multiplying the major and minor axis of the tumour as measured with a vernier calliper.

### Statistical analysis

Different morphometric parameters were tested for significance using the Mann-Whitney test. Significance test for the *in-silico* data was performed using either Mann-Whitney or Kruskal-Wallis test. The Mann–Whitney U-test was used to analyse the statistical significance of relative Golgi area changes and relative fluorescence intensity in the 3D culture immunostaining. The different parameters for analysing cell migration was tested for significance using the Mann-Whitney test. Statistical analysis for mice tumour area was performed using the 2-way ANOVA followed by Sidak’s multiple comparison test. P<0.05 was considered statistically significant. Graph Pad Prism software (Graph Pad Software, La Jolla, CA, USA) was used to analyse data.

### Ethics approvals

Ethics approval from Institutional Human Ethics Committee (IHEC) was obtained for using patient paraffin embedded tissue blocks in this study (IHEC/Admin/2021/012). Written informed consent was obtained by Prashanti Cancer Care Mission, Pune from all patients, and the study was conducted in accordance with the Declaration of Helsinki, institutional guidelines, and all local, state and national regulations. For preparing Api5 OE MCF10A cells, Api5 KD MCF10AT1 and Api5 KD MCF10CA1a cells Institutional Biosafety Committee (IBSC) clearance was approved by the institute. The *in vivo* tumorigenicity studies were approved by the Institutional Animal Ethics Committee (IAEC) (IAEC/2018_02/010 and IISER_Pune/IAEC/2021_01/06).

## Acknowledgements

We thank Dr Richa Rikhy, (IISER Pune, India) and Dr Sorab Dalal (ACTREC, Mumbai, India) for their useful suggestions. The authors sincerely acknowledge Dr Girish Deshpande (Princeton University, USA) and Prof L S Shashidhara (Ashoka University, India) for their valuable comments and edits in the manuscript. We also thank Dr Manas Kumar Santra (NCCS, Pune, India) and IISER Pune-BD FACS facility for help with sorting of cells. The authors acknowledge Dr T.S. Sridhar and his lab members (SJRI, Bangalore, India) for help with IHC protocols and Dr C.B. Koppikar and team (Prashanti Cancer Care Mission, Pune, India) for providing the paraffin-embedded tissue blocks. We thank Prof. Anjan Banerjee for allowing us to use the microtome for tissue sectioning. We also would like to acknowledge the IISER Pune Microscopy Facility for access to equipment and infrastructure and National Facility for Gene Function in Health and Disease (IISER, Pune, India) for access to the animal facility and support for experiments. We also thank Lahiri lab members for helpful comments and discussions.

## Author contributions

Conceptualisation: A.K, M.L.; Methodology: A.K, D.P., R.M., G.G., M.L.; Validation: A.K; Formal analysis: A.K, D.P., R.M., G.G., M.L.; Investigation: A.K, D.P., R.M., G.G.; Writing - original draft: A.K.; Writing - review & editing: M.L.; Visualisation: A.K, M.L.; Supervision: M.L.; Project administration: M.L.; Funding acquisition: M.L.

## References

Aiello NM, Kang Y. 2019. Context-dependent EMT programs in cancer metastasis. J Exp Med 216: 1016–1026.

Almasan A, Ashkenazi A. 2003. Apo2L/TRAIL: apoptosis signaling, biology, and potential for cancer therapy. Cytokine Growth Factor Rev 14: 337–348.

Anandi L, Chakravarty V, Ashiq KA, Bodakuntla S, Lahiri M. 2017. DNA-dependent protein kinase plays a central role in transformation of breast epithelial cells following alkylation damage. J Cell Sci 130: 3749–3763.

Anandi VL, Ashiq KA, Nitheesh K, Lahiri M. 2016. Platelet-activating factor promotes motility in breast cancer cells and disrupts non-transformed breast acinar structures. Oncol Rep 35: 179–188.

Basset C, Bonnet-Magnaval F, Navarro MG, Touriol C, Courtade M, Prats H, Garmy-Susini B, Lacazette E. 2017. Api5 a new cofactor of estrogen receptor alpha involved in breast cancer outcome. Oncotarget 8: 52511–52526.

Bessette DC, Tilch E, Seidens T, Quinn MC, Wiegmans AP, Shi W, Cocciardi S, McCart-Reed A, Saunus JM, Simpson PT et al. 2015. Using the MCF10A/MCF10CA1a Breast Cancer Progression Cell Line Model to Investigate the Effect of Active, Mutant Forms of EGFR in Breast Cancer Development and Treatment Using Gefitinib. PLoS One 10: e0125232.

Blockhuys S, Zhang X, Wittung-Stafshede P. 2020. Single-cell tracking demonstrates copper chaperone Atox1 to be required for breast cancer cell migration. Proc Natl Acad Sci U S A 117: 2014–2019.

Bodakuntla S, Libi AV, Sural S, Trivedi P, Lahiri M. 2014. N-nitroso-N-ethylurea activates DNA damage surveillance pathways and induces transformation in mammalian cells. BMC Cancer 14: 287.

Bong SM, Bae SH, Song B, Gwak H, Yang SW, Kim S, Nam S, Rajalingam K, Oh SJ, Kim TW et al. 2020. Regulation of mRNA export through API5 and nuclear FGF2 interaction. Nucleic Acids Res 48: 6340–6352.

Bousquet G, Feugeas JP, Gu Y, Leboeuf C, Bouchtaoui ME, Lu H, Espie M, Janin A, Benedetto MD. 2019. High expression of apoptosis protein (Api-5) in chemoresistant triple-negative breast cancers: an innovative target. Oncotarget 10: 6577–6588.

Cho H, Chung JY, Song KH, Noh KH, Kim BW, Chung EJ, Ylaya K, Kim JH, Kim TW, Hewitt SM et al. 2014. Apoptosis inhibitor-5 overexpression is associated with tumor progression and poor prognosis in patients with cervical cancer. BMC Cancer 14: 545.

Cho KB, Cho MK, Lee WY, Kang KW. 2010. Overexpression of c-myc induces epithelial mesenchymal transition in mammary epithelial cells. Cancer Lett 293: 230–239.

Dawson PJ, Wolman SR, Tait L, Heppner GH, Miller FR. 1996. MCF10AT: a model for the evolution of cancer from proliferative breast disease. Am J Pathol 148: 313–319.

Debnath J, Brugge JS. 2005. Modelling glandular epithelial cancers in three-dimensional cultures. Nat Rev Cancer 5: 675–688.

Debnath J, Muthuswamy SK, Brugge JS. 2003a. Morphogenesis and oncogenesis of MCF-10A mammary epithelial acini grown in three-dimensional basement membrane cultures. Methods 30: 256–268.

Debnath J, Walker SJ, Brugge JS. 2003b. Akt activation disrupts mammary acinar architecture and enhances proliferation in an mTOR-dependent manner. J Cell Biol 163: 315–326.

DuChez BJ. 2017. Automated Tracking of Cell Migration with Rapid Data Analysis. Curr Protoc Cell Biol 76: 12 12 11–12 12 16.

Duffy MJ, Synnott NC, Crown J. 2018. Mutant p53 in breast cancer: potential as a therapeutic target and biomarker. Breast Cancer Res Treat 170: 213–219.

Gaiko-Shcherbak A, Fabris G, Dreissen G, Merkel R, Hoffmann B, Noetzel E. 2015. The Acinar Cage: Basement Membranes Determine Molecule Exchange and Mechanical Stability of Human Breast Cell Acini. PLoS One 10: e0145174.

Garcia-Jove Navarro M, Basset C, Arcondeguy T, Touriol C, Perez G, Prats H, Lacazette E. 2013. Api5 contributes to E2F1 control of the G1/S cell cycle phase transition. PLoS One 8: e71443.

Green AR, Aleskandarany MA, Agarwal D, Elsheikh S, Nolan CC, Diez-Rodriguez M, Macmillan RD, Ball GR, Caldas C, Madhusudan S et al. 2016. MYC functions are specific in biological subtypes of breast cancer and confers resistance to endocrine therapy in luminal tumours. Br J Cancer 114: 917–928.

Guiu S, Michiels S, Andre F, Cortes J, Denkert C, Di Leo A, Hennessy BT, Sorlie T, Sotiriou C, Turner N et al. 2012. Molecular subclasses of breast cancer: how do we define them? The IMPAKT 2012 Working Group Statement. Ann Oncol 23: 2997–3006.

Halaoui R, McCaffrey L. 2015. Rewiring cell polarity signaling in cancer. Oncogene 34: 939–950.

Halaoui R, Rejon C, Chatterjee SJ, Szymborski J, Meterissian S, Muller WJ, Omeroglu A, McCaffrey L. 2017. Progressive polarity loss and luminal collapse disrupt tissue organisation in carcinoma. Genes Dev 31: 1573–1587.

Han BG, Kim KH, Lee SJ, Jeong KC, Cho JW, Noh KH, Kim TW, Kim SJ, Yoon HJ, Suh SW et al. 2012. Helical repeat structure of apoptosis inhibitor 5 reveals protein-protein interaction modules. J Biol Chem 287: 10727–10737.

Hanahan D, Weinberg RA. 2011. Hallmarks of cancer: the next generation. Cell 144: 646–674.

Imbalzano KM, Tatarkova I, Imbalzano AN, Nickerson JA. 2009. Increasingly transformed MCF-10A cells have a progressively tumor-like phenotype in three-dimensional basement membrane culture. Cancer Cell Int 9: 7.

Imre G, Berthelet J, Heering J, Kehrloesser S, Melzer IM, Lee BI, Thiede B, Dotsch V, Rajalingam K. 2017. Apoptosis inhibitor 5 is an endogenous inhibitor of caspase-2. EMBO Rep 18: 733–744.

Jagot-Lacoussiere L, Kotula E, Villoutreix BO, Bruzzoni-Giovanelli H, Poyet JL. 2016. A Cell-Penetrating Peptide Targeting AAC-11 Specifically Induces Cancer Cells Death. Cancer Res 76: 5479–5490.

Jang HS, Woo SR, Song KH, Cho H, Chay DB, Hong SO, Lee HJ, Oh SJ, Chung JY, Kim JH et al. 2017. API5 induces cisplatin resistance through FGFR signaling in human cancer cells. Exp Mol Med 49: e374.

Jansen MP, Foekens JA, van Staveren IL, Dirkzwager-Kiel MM, Ritstier K, Look MP, Meijer-van Gelder ME, Sieuwerts AM, Portengen H, Dorssers LC et al. 2005. Molecular classification of tamoxifen-resistant breast carcinomas by gene expression profiling. J Clin Oncol 23: 732–740.

Javed A, Lteif A. 2013. Development of the human breast. Semin Plast Surg 27: 5–12.

Javier RT. 2008. Cell polarity proteins: common targets for tumorigenic human viruses. Oncogene 27: 7031–7046.

Jolly MK, Boareto M, Huang B, Jia D, Lu M, Ben-Jacob E, Onuchic JN, Levine H. 2015. Implications of the Hybrid Epithelial/Mesenchymal Phenotype in Metastasis. Front Oncol 5: 155.

Kim HD, Guo TW, Wu AP, Wells A, Gertler FB, Lauffenburger DA. 2008. Epidermal growth factor-induced enhancement of glioblastoma cell migration in 3D arises from an intrinsic increase in speed but an extrinsic matrix- and proteolysis-dependent increase in persistence. Mol Biol Cell 19: 4249–4259.

Kim JW, Cho HS, Kim JH, Hur SY, Kim TE, Lee JM, Kim IK, Namkoong SE. 2000. AAC-11 overexpression induces invasion and protects cervical cancer cells from apoptosis. Lab Invest 80: 587–594.

Krejci P, Pejchalova K, Rosenbloom BE, Rosenfelt FP, Tran EL, Laurell H, Wilcox WR. 2007. The antiapoptotic protein Api5 and its partner, high molecular weight FGF2, are up-regulated in B cell chronic lymphoid leukemia. J Leukoc Biol 82: 1363–1364.

Mailleux AA, Overholtzer M, Schmelzle T, Bouillet P, Strasser A, Brugge JS. 2007. BIM regulates apoptosis during mammary ductal morphogenesis, and its absence reveals alternative cell death mechanisms. Dev Cell 12: 221–234.

Millarte V, Farhan H. 2012. The Golgi in cell migration: regulation by signal transduction and its implications for cancer cell metastasis. ScientificWorldJournal 2012: 498278.

Morris EJ, Michaud WA, Ji JY, Moon NS, Rocco JW, Dyson NJ. 2006. Functional identification of Api5 as a suppressor of E2F-dependent apoptosis in vivo. PLoS Genet 2: e196.

Mossahebi-Mohammadi M, Quan M, Zhang JS, Li X. 2020. FGF Signaling Pathway: A Key Regulator of Stem Cell Pluripotency. Front Cell Dev Biol 8: 79.

Nagy A, Munkacsy G, Gyorffy B. 2021. Pancancer survival analysis of cancer hallmark genes. Sci Rep 11: 6047.

Noh KH, Kim SH, Kim JH, Song KH, Lee YH, Kang TH, Han HD, Sood AK, Ng J, Kim K et al. 2014. API5 confers tumoral immune escape through FGF2-dependent cell survival pathway. Cancer Res 74: 3556–3566.

O’Reilly LA, Kruse EA, Puthalakath H, Kelly PN, Kaufmann T, Huang DC, Strasser A. 2009. MEK/ERK-mediated phosphorylation of Bim is required to ensure survival of T and B lymphocytes during mitogenic stimulation. J Immunol 183: 261–269.

Okada T, Enkhjargal B, Travis ZD, Ocak U, Tang J, Suzuki H, Zhang JH. 2019. FGF-2 Attenuates Neuronal Apoptosis via FGFR3/PI3k/Akt Signaling Pathway After Subarachnoid Hemorrhage. Mol Neurobiol 56: 8203–8219.

Park SJ, Yoon BH, Kim SK, Kim SY. 2019. GENT2: an updated gene expression database for normal and tumor tissues. BMC Med Genomics 12: 101.

Partanen JI, Nieminen AI, Makela TP, Klefstrom J. 2007. Suppression of oncogenic properties of c-Myc by LKB1-controlled epithelial organisation. Proc Natl Acad Sci U S A 104: 14694–14699.

Paul CD, Mistriotis P, Konstantopoulos K. 2017. Cancer cell motility: lessons from migration in confined spaces. Nat Rev Cancer 17: 131–140.

Petrosyan A. 2015. Onco-Golgi: Is Fragmentation a Gate to Cancer Progression? Biochem Mol Biol J 1.

Plati J, Bucur O, Khosravi-Far R. 2008. Dysregulation of apoptotic signaling in cancer: molecular mechanisms and therapeutic opportunities. J Cell Biochem 104: 1124–1149.

Plati J, Bucur O, Khosravi-Far R. 2011. Apoptotic cell signaling in cancer progression and therapy. Integr Biol (Camb) 3: 279–296.

Pouthas F, Girard P, Lecaudey V, Ly TB, Gilmour D, Boulin C, Pepperkok R, Reynaud EG. 2008. In migrating cells, the Golgi complex and the position of the centrosome depend on geometrical constraints of the substratum. J Cell Sci 121: 2406–2414.

Qu Y, Han B, Yu Y, Yao W, Bose S, Karlan BY, Giuliano AE, Cui X. 2015. Evaluation of MCF10A as a Reliable Model for Normal Human Mammary Epithelial Cells. PLoS One 10: e0131285.

Ramdas P, Rajihuzzaman M, Veerasenan SD, Selvaduray KR, Nesaretnam K, Radhakrishnan AK. 2011. Tocotrienol-treated MCF-7 human breast cancer cells show down-regulation of API5 and up-regulation of MIG6 genes. Cancer Genomics Proteomics 8: 19–31.

Reginato MJ, Mills KR, Becker EB, Lynch DK, Bonni A, Muthuswamy SK, Brugge JS. 2005. Bim regulation of lumen formation in cultured mammary epithelial acini is targeted by oncogenes. Mol Cell Biol 25: 4591–4601.

Roche J. 2018. The Epithelial-to-Mesenchymal Transition in Cancer. Cancers (Basel) 10.

Saitoh M. 2018. Involvement of partial EMT in cancer progression. J Biochem 164: 257–264.

Santner SJ, Dawson PJ, Tait L, Soule HD, Eliason J, Mohamed AN, Wolman SR, Heppner GH, Miller FR. 2001. Malignant MCF10CA1 cell lines derived from premalignant human breast epithelial MCF10AT cells. Breast Cancer Res Treat 65: 101–110.

Saraste J, Prydz K. 2019. A New Look at the Functional Organization of the Golgi Ribbon. Front Cell Dev Biol 7: 171.

Sasaki H, Moriyama S, Yukiue H, Kobayashi Y, Nakashima Y, Kaji M, Fukai I, Kiriyama M, Yamakawa Y, Fujii Y. 2001. Expression of the antiapoptosis gene, AAC-11, as a prognosis marker in non-small cell lung cancer. Lung Cancer 34: 53–57.

Shahbandi A, Nguyen HD, Jackson JG. 2020. TP53 Mutations and Outcomes in Breast Cancer: Reading beyond the Headlines. Trends Cancer 6: 98–110.

Sharma VK, Lahiri M. 2021. Interplay between p300 and HDAC1 regulate acetylation and stability of Api5 to regulate cell proliferation. Sci Rep 11: 16427.

Simpson DR, Yu M, Zheng S, Zhao Z, Muthuswamy SK, Tansey WP. 2011. Epithelial cell organisation suppresses Myc function by attenuating Myc expression. Cancer Res 71: 3822–3830.

Song KH, Cho H, Kim S, Lee HJ, Oh SJ, Woo SR, Hong SO, Jang HS, Noh KH, Choi CH et al. 2017. API5 confers cancer stem cell-like properties through the FGF2-NANOG axis. Oncogenesis 6: e285.

Soule HD, Maloney TM, Wolman SR, Peterson WD, Jr., Brenz R, McGrath CM, Russo J, Pauley RJ, Jones RF, Brooks SC. 1990. Isolation and characterisation of a spontaneously immortalised human breast epithelial cell line, MCF-10. Cancer Res 50: 6075–6086.

Swamydas M, Eddy JM, Burg KJ, Dreau D. 2010. Matrix compositions and the development of breast acini and ducts in 3D cultures. In Vitro Cell Dev Biol Anim 46: 673–684.

Tang CX, Luan L, Zhang L, Wang Y, Liu XF, Wang J, Xiong Y, Wang D, Huang LY, Gao DS. 2019. Golgin-160 and GMAP210 play an important role in U251 cells migration and invasion initiated by GDNF. PLoS One 14: e0211501.

Tewari M, Yu M, Ross B, Dean C, Giordano A, Rubin R. 1997. AAC-11, a novel cDNA that inhibits apoptosis after growth factor withdrawal. Cancer Res 57: 4063–4069.

Wang M, Li J, Huang J, Luo M. 2020. The Predictive Role of PIK3CA Mutation Status on PI3K Inhibitors in HR+ Breast Cancer Therapy: A Systematic Review and Meta-Analysis. Biomed Res Int 2020: 1598037.

Wong RS. 2011. Apoptosis in cancer: from pathogenesis to treatment. J Exp Clin Cancer Res 30: 87.

Xu S, Feng Y, Zhao S. 2019. Proteins with Evolutionarily Hypervariable Domains are Associated with Immune Response and Better Survival of Basal-like Breast Cancer Patients. Comput Struct Biotechnol J 17: 430–440.

Yin S, Cheryan VT, Xu L, Rishi AK, Reddy KB. 2017. Myc mediates cancer stem-like cells and EMT changes in triple negative breast cancers cells. PLoS One 12: e0183578.

