## Supplementary Information for "Altered expression of Api5 affects breast carcinogenesis by modulating FGF2 signalling"

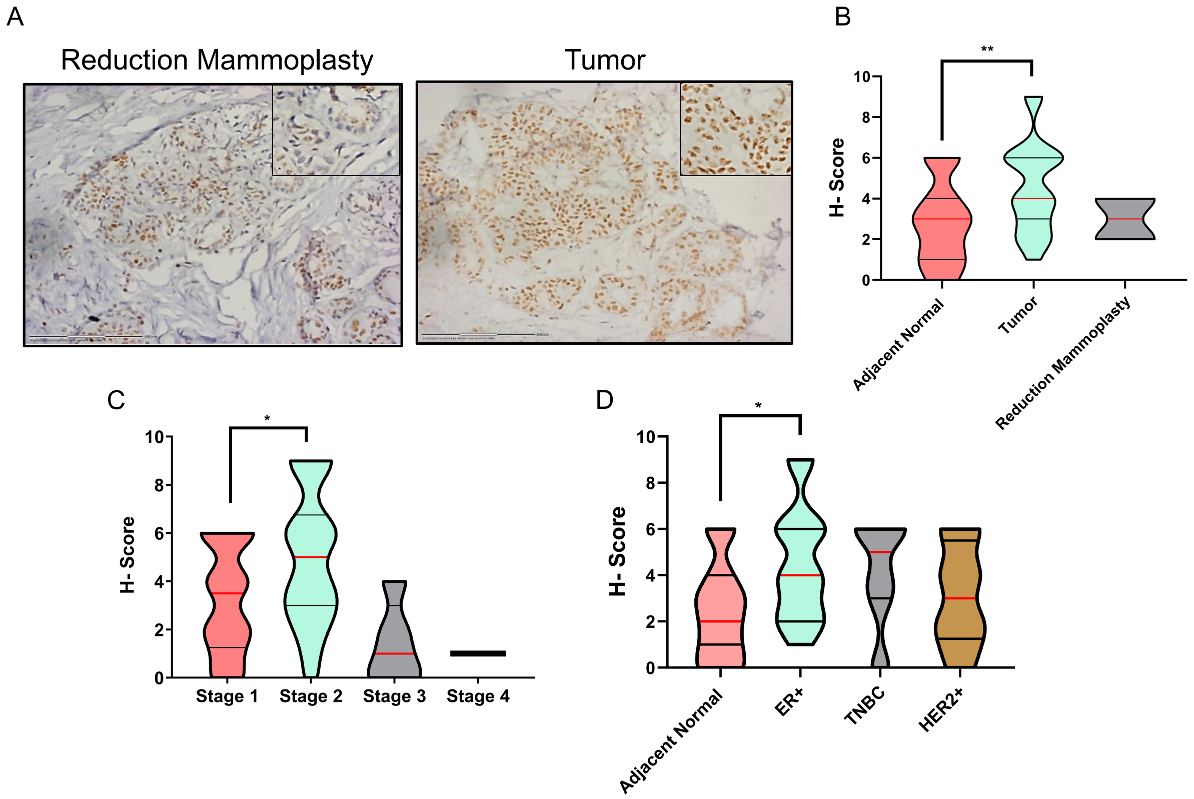


**Supplementary Figure 1:** (A) Representative image of Api5 immunostained section of a paraffin-embedded breast cancer tissue block and the corresponding reduction mammoplasty section. Inset shows a zoomed-in image. H score of tissue sample compared between (B) adjacent normal and tumour tissues, (C) different stages of breast cancer and (D) subtypes of breast cancer plotted as violin plots. Statistical analysis was performed using the Mann-Whitney test. *P<0.05, **P<0.01, ***P<0.001and ****P<0.0001. N≥25.

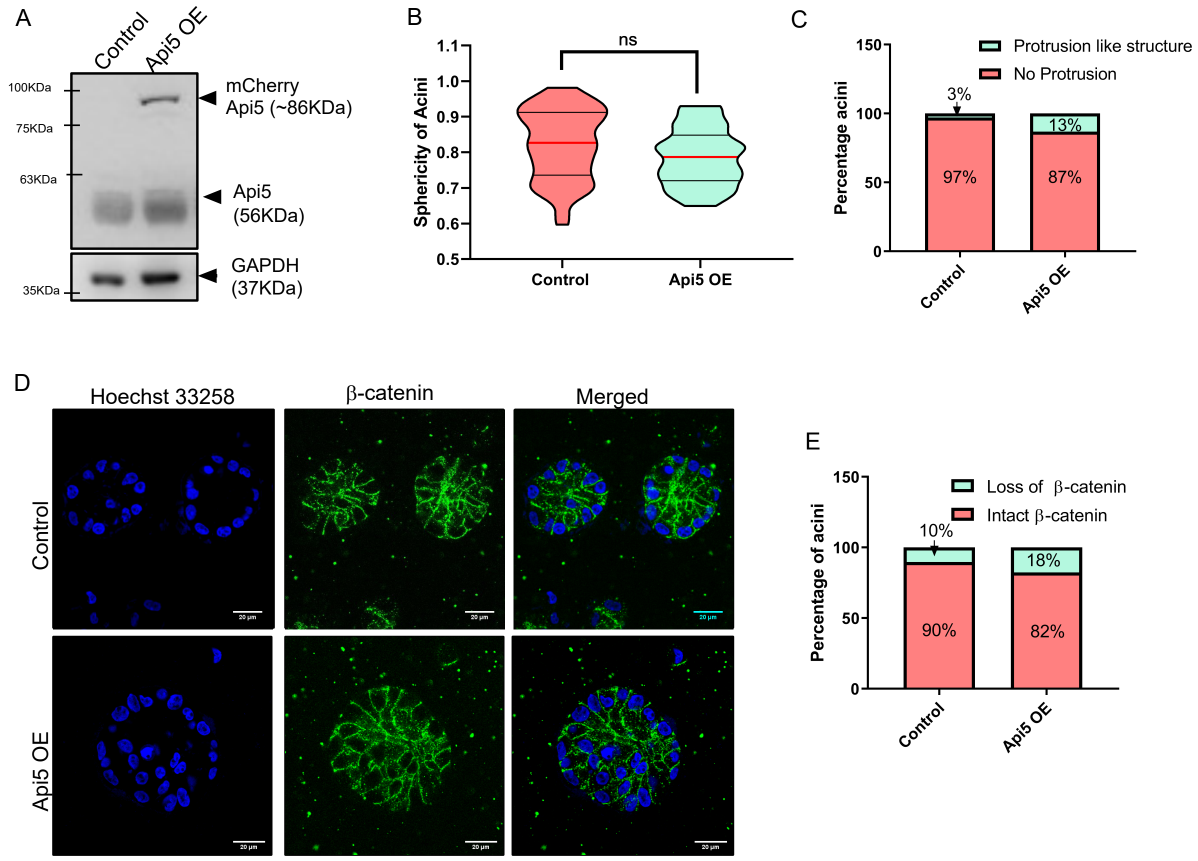


**Supplementary Figure 2:** (A) Western blot image showing overexpression of Api5 in MCF10A lysates collected from day 16 acini. (B) Graph showing sphericity of day 16 acini analysed based on phalloidin staining using Huygens software (C) Graph showing percentage of acini with protrusion-like structure manually analysed based on phalloidin staining in MCF10A cells. (D) Representative image of β-catenin staining (green) in day 16 acini. (E) Graph showing percentage of acini with intact and loss of β-catenin.

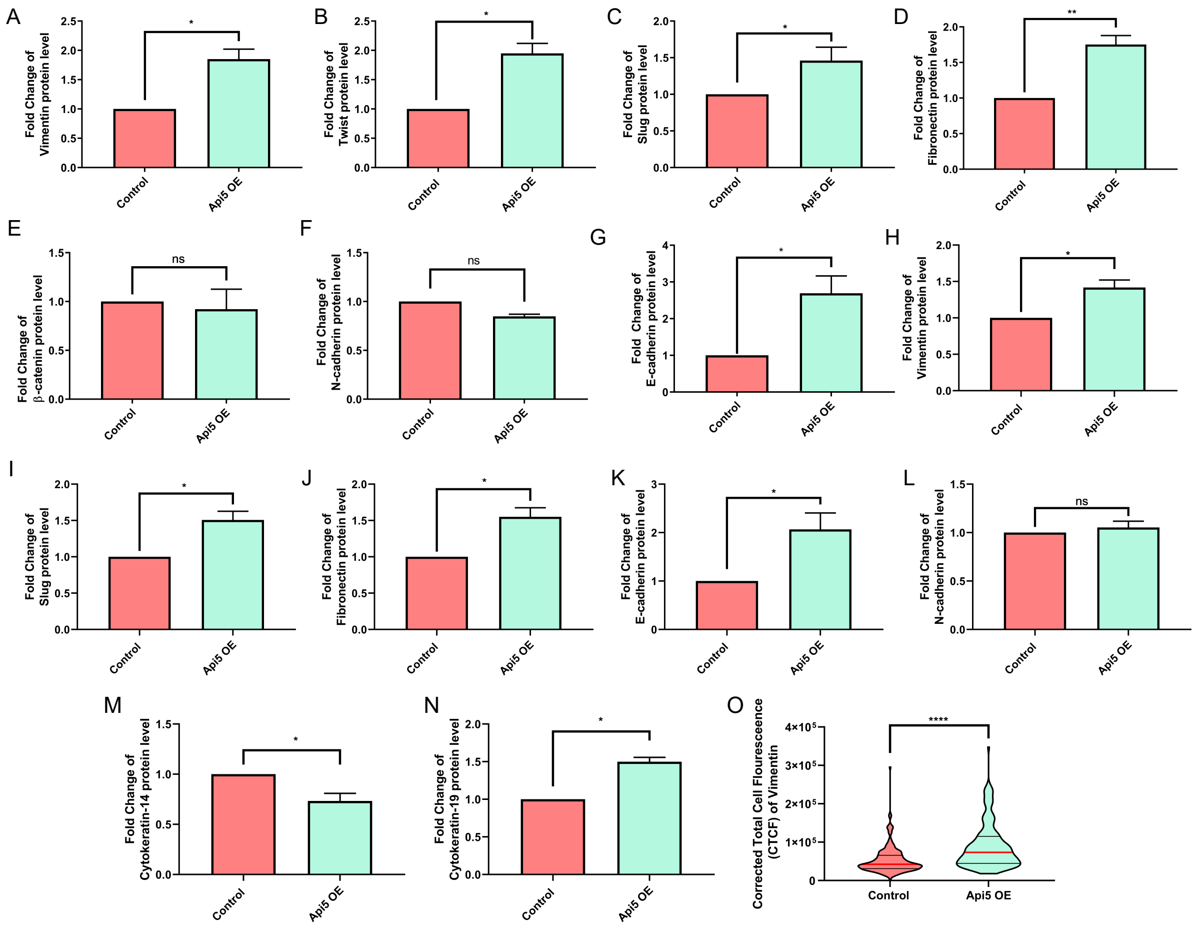


**Supplementary Figure 3:** Breast epithelial cells overexpressing Api5 acquires partial EMT-like characteristics. Graph showing fold change of (A) Vimentin, (B) Twist (C) Slug, (D) Fibronectin, (E) β-catenin, (F) N-cadherin and (G) E-cadherin protein expression lysates collected from day 16 lysates of 3D culture. Graph showing fold change of (H) Vimentin, (I) Slug, (J) Fibronectin, (K) E-cadherin, (L) N-cadherin, (M) Cytokeratin-14 and (N) Cytokeratin-19 in lysates collected from 3D dissociated cells of control and Api5 OE MCF10A. (O) Violin plot showing the corrected fluorescence intensity of vimentin in 3D dissociated cells. Fold change in protein levels were calculated in comparison to the control lane and normalised to GAPDH. Statistical analysis was performed using the paired t-test. *P<0.05, **P<0.01, ***P<0.001and ****P<0.0001. Data pooled from N≥3 independent experiments.

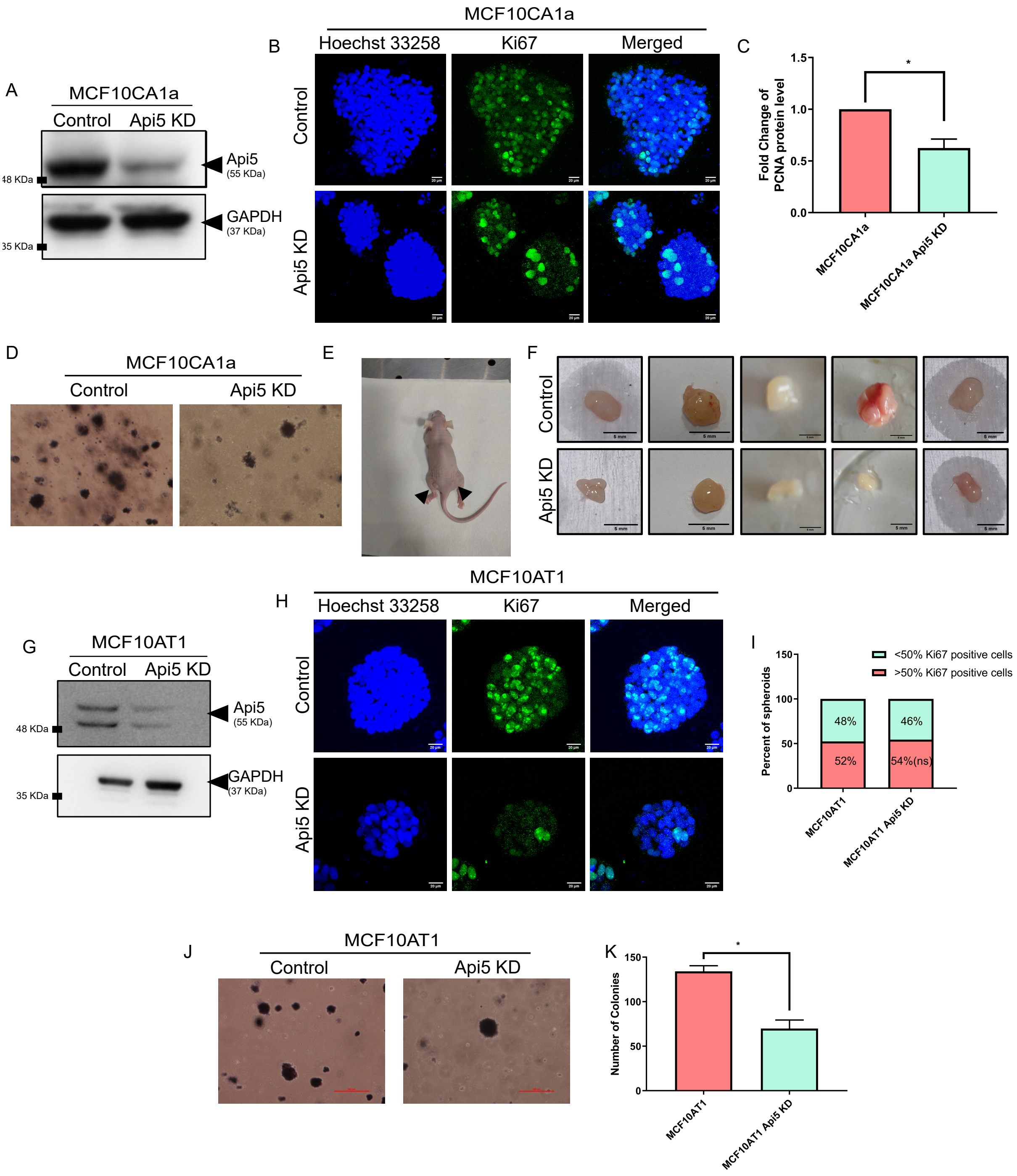


**Supplementary Figure 4** (A) Western blot showing Api5 levels in stable cells of MCF10CA1a transduced with shApi5 construct. (B) Representative image of Api5 KD MCF10CA1a cells immunostained for proliferation marker, Ki67 after 7 days of culturing in Matrigel® (Ki67: green, nuclei: blue) (C) Graph showing the fold change in PCNA protein levels in lysates collected from day 7 spheroid cultures. (D) Representative image showing colony formed in soft agar assay by MCF10CA1a cells with Api5 KD stained with MTT after 21 days of seeding. (E) Representative image of mice sacrificed for tumour dissection post 8 weeks. Left flank was injected with control cells and the right flank with Api5 KD cells. (F) Representative images of tumour dissected from flanks of athymic mice after week 8 post subcutaneous injection. The upper panel shows MCF10CA1a control cells, while the lower panel shows Api5 KD MCF10CA1a. (G) Western blot showing Api5 levels in stable cells of MCF10AT1 transduced with shApi5 construct. (H) Representative image of Api5 KD MCF10AT1 cells immunostained for Ki67 after 7 days of culturing in Matrigel® (Ki67: green, nuclei: blue). (I) Graph showing the percentage of spheroids with 50% or more cells showing Ki67 staining. (J) Representative image showing colony formed in soft agar assay by MCF10AT1 cells with Api5 KD stained with MTT after 21 days of seeding. (K) Bar graph representing the number of colonies formed by MCF10AT1 cells compared between control and Api5 KD. Statistical analysis was performed using the Mann-Whitney test. *P<0.05, **P<0.01, ***P<0.001and ****P<0.0001 or paired t-test. *P<0.05, **P<0.01, ***P<0.001and ****P<0.0001. Data pooled from n>4 independent.

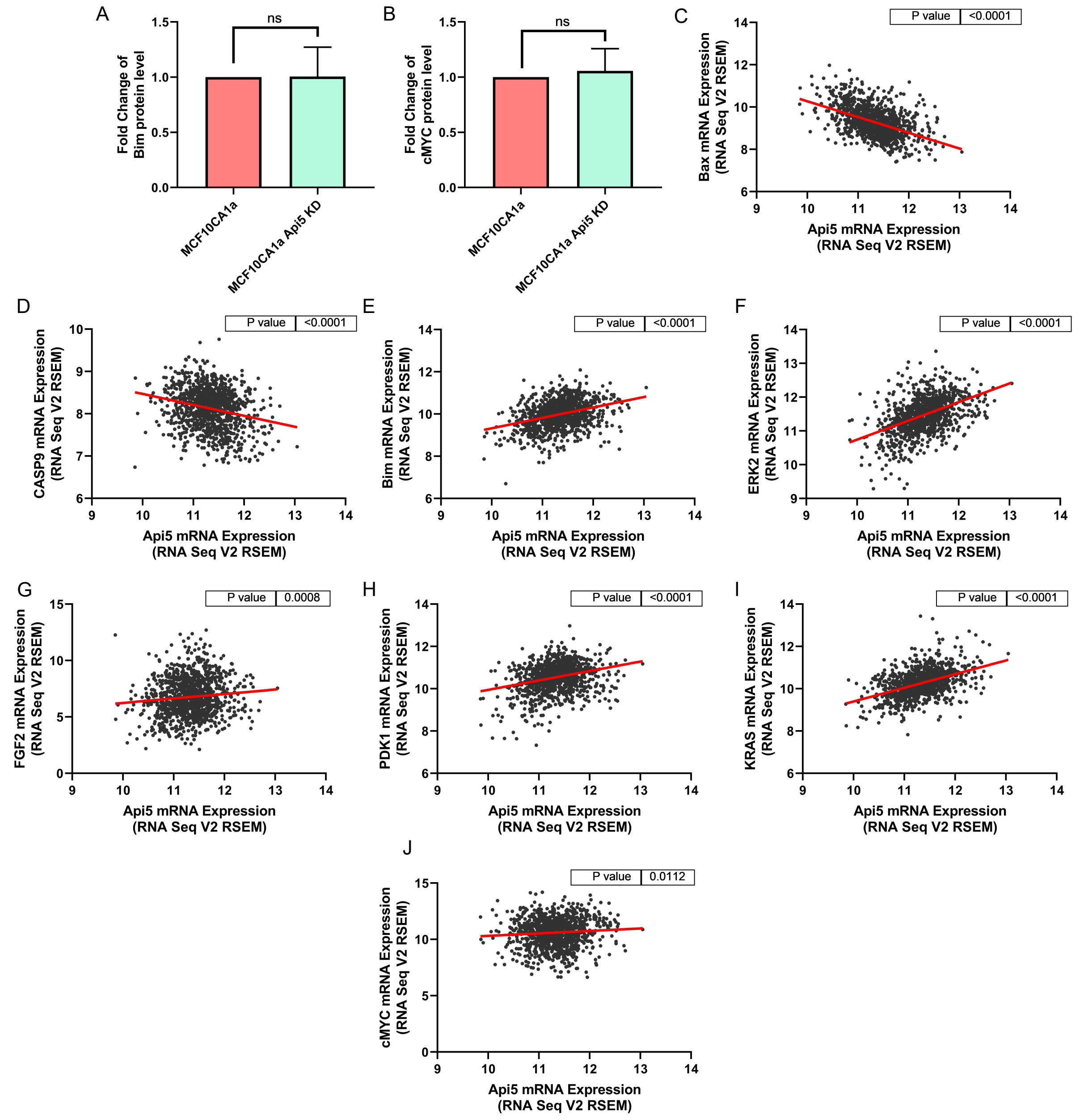


**Supplementary Figure 5:** Bar diagram representing the expression of (A) Bim and (B) cMYC in lysates collected from day 7 MCF10CA1a spheroids. (C-J) TCGA-based co-expression analysis performed between transcript expression of proteins involved in the pathway depicted in Figure 7 and Api5. Graphs are generated using TCGA data downloaded using UCSC Xena Browser(Goldman et al. 2020).
